# CRUMBLER: A Tool for the Prediction of Ancestry in Cattle

**DOI:** 10.1101/396341

**Authors:** Tamar E. Crum, Robert D. Schnabel, Jared E. Decker, Luciana CA Regitano, Jeremy F. Taylor

## Abstract

**Background:** In many beef and some dairy production systems, crossbreeding is used to take advantage of breed complementarity and heterosis. Admixed animals are frequently identified by their coat color and body conformation phenotypes, however, without pedigree information it is not possible to identify the expected breed composition of an admixed animal and in the presence of selection, the actual composition may differ from expectation. As the roles of DNA and genotype data become more pervasive in animal agriculture, a systematic method for estimating the breed composition (the proportions of an animal’s genome originating from ancestral pure breeds) has utility for a variety of downstream analyses including the estimation of genomic breeding values for crossbred animals, the estimation of quantitative trait locus effects, and heterosis and heterosis retention in advanced generation composite animals. Currently, there is no automated or semi-automated ancestry estimation platform for cattle and the objective of this study was to evaluate the utility of extant public software for ancestry estimation and determine the effects of reference population size and composition and number of utilized single nucleotide polymorphism loci on ancestry estimation. We also sought to develop an analysis pipeline that would simplify this process for members of the livestock genomics research community.

**Results:** We developed and tested a tool, “CRUMBLER”, to estimate the global ancestry of cattle using ADMIXTURE and SNPweights based on a defined reference panel.

CRUMBLER, was developed and evaluated in cattle, but is a species agnostic pipeline that facilitates the streamlined estimation of breed composition for individuals with potentially complex ancestries using publicly available global ancestry software and a specified reference population SNP dataset. We developed the reference panel from a large cattle genotype data set and breed association pedigree information using iterative analyses to identify purebred individuals that were representative of each breed. We also evaluated the numbers of markers necessary for breed composition estimation and simulated genotypes for advanced generation composite animals to evaluate the precision of the developed tool.

**Conclusion:** The developed CRUMBLER pipeline extracts a specified subset of genotypes that is common to all current commercially available genotyping platforms, processes these into the file formats required for the analysis software, and predicts admixture proportions using the specified reference population allele frequencies.

## Background

Estimation of the breed composition of individuals with complex ancestries has utility for estimating breed direct and heterosis effects as well as for the estimation of the additive genetic merit of these individuals. It also has value for identifying the breed composition of training populations used for genomic selection and hence the identification of target breeds in which the developed prediction equations may have some relevance. Visual classification of cattle based on breed characteristics suffers from similar problems as the self-identification of ethnicity in humans [1], as most breed characteristics are determined by alleles at relatively few loci. For example, recent extensive crossing with Angus cattle in the U.S. produces a black hided animal which masks all other solid coat colors found in other breeds and requires only a single dominant allele at the *MC1R* locus. As a consequence, black-hided cattle have a “cryptic” population structure [1,2] and the visual classification of black-hided animals for branded beef programs can result in the marketing of animals with vastly different Angus genome content.

In the U.S. and many other countries, the breed of an animal is associated with its being registered with a breed association which requires that both parents of the animal be identified and also registered with the association. For the previous 50 years, parentage has been validated by each breed association using blood or, more recently, DNA typing. Many breed associations have closed herdbooks which means, in theory, that the pedigrees of all animals can be traced back to the animals that founded the breed’s herdbook. Other breed associations have open herdbooks, which means that crossbred animals can be registered with the breed if they have been graded up by crossbreeding to purebred status with the expectation that a certain percentage of their genome (e.g., 15/16ths) originates from the respective breed based upon pedigree records and parentage validation. Pedigree errors that occurred prior to, or that were not identified following the implementation of blood typing and DNA testing, lead to admixed animals being incorrectly classified as fullblood and incorrectly identified admixture proportions in purebred animals. The effects of recombination, random assortment of chromosomes into gametes and selection can also lead to considerable variation in the extent of identity by descent between relatives separated by more than a single meiosis and can also lead to admixture proportions that differ substantially from expectation based on pedigree.

Crossbreeding is extensively used in commercial beef production and in other livestock species production systems to capitalize on the effects of breed complementarity and heterosis resulting in herds of females that may have very complex ancestries that frequently use fullblood or purebred bulls sourced from registered breeders. Changes in the decision as to which breed of bull to use can result in large changes in admixture proportions of replacement cows and marketed steers between years and large differences can occur between herds for the same reason. When commercially sourced animals are used to generate resource populations to study the genomics of economically important traits such as feed conversion efficiency [3,4] or bovine respiratory disease [5], the presence of extensive admixture in the phenotyped and genotyped animals may impact the GWAA [3,4] and leads to the training of genomic prediction models in populations for which the breed composition is not understood. As a consequence, the utility of these models in other industry populations, including the registered breeds in which the majority of genetic improvement is generated is also not understood.

As the number of genotyped beef animals has increased, the need to classify the breed composition of these animals has necessitated the development of a precise and accurate method for estimating breed composition in cattle based on single nucleotide polymorphism (SNP) data. Iterative ancestry estimation analyses performed using different software input parameters may identify those that cause output sensitivity and can lead to an interpretation of population structure that is close to the truth [6]. We developed the CRUMBLER analysis pipeline to streamline the genomic estimation of breed composition of crossbred cattle using high-density SNP genotype data, publicly available software, and a reference panel containing genotypes for members of cattle breeds that are numerically important in North America. The CRUMBLER pipeline is species agnostic and could be adapted for breed composition estimation in other species. CRUMBLER and the reference panel data are available on GitHub (https://github.com/tamarcrum/CRUMBLER). This pipeline tool is released under the GNU General Public License.

## Materials and Methods

### Genotype data

From among the numerically most important cattle breeds in North America, in terms of their annual numbers of animal registrations, a list was compiled to define the target breeds for reference panel development. Composite breeds, such as Brangus and Braford, were not included in this list due to lack of available genotype data, but the progenitor Angus, Hereford and Brahman breeds were included. Breeds such as N’Dama, representing African taurine, and Nelore and Brahman, representing *Bos taurus indicus* cattle, were included. We also initially included breeds that were likely to be involved in early crossbreeding of cattle in the U.S. (Texas Longhorn).

From the 170,544 cattle with high-density SNP genotypes stored within the University of Missouri Animal Genomics genotype database, we extracted genotypes for 48,776 animals identified as being registered with one of the numerically important U.S. Breed Associations or belonging to other world breeds. Pedigree data were also obtained for these animals from each of the Breed Associations, where available (Table 1). These individuals had been genotyped using at least one of 9 different genotyping platforms currently used internationally to genotype cattle including the GeneSeek (Lincoln, NE) GGP-90KT, GGP-F250, GGP-HDV3, GGP-LDV1, GGP-LDV3, and GGP-LDV4 assays, the Illumina (San Diego, CA) BovineHD and BovineSNP50 assays, and the Zoetis (Kalamazoo, MI) i50K assay. The numbers of variants queried by each assay and the number of individuals genotyped using each platform are shown in Table 2.

**Table 1.** Genotype data for 48,776 registered individuals from 20 breeds were used to establish the reference population.

| **Breed** | **No. Registered Individuals** | **No. FullBlood Individuals<sup>a</sup>** | **No. Individuals Assigned to Breed<sup>b</sup>** | **Sampled Individuals<sup>c</sup>** | **No. Individual After Pedigree and SNPweights<sup>d</sup>** |
| --- | --- | --- | --- | --- | --- |
| Angus | 5552 | 5552 | 485 | 200 | 200 |
| Hereford | 969 | 969 | 348 | 200 | 200 |
| Limousin | 2734 | 321 | 367 | 200 | 200 |
| Charolais | 1542 | 1489 | 1542 | 200 | 200 |
| Simmental | 15858 | 337 | 1583 | 200 | 196 |
| Japanese Black | 97 | 97 | 97 | 97 | 94 |
| Braunvieh | 148 | 69 | 148 | 148 | 69 |
| Gelbvieh | 12835 | 51 | 6000 | 200 | 51 |
| Romagnola | 37 | 37 | 37 | 37 | 37 |
| Salers | 68 | 68 | 0 | 0 | 0 |
| Texas Longhorn | 45 | 45 | 45 | 0 | 0 |
| Shorthorn | 291 | 178 | 166 | 166 | 178 |
| Red Angus | 1377 | 1377 | 124 | 124 | 124 |
| Holstein | 5816 | 5816 | 5816 | 200 | 197 |
| Jersey | 119 | 119 | 119 | 119 | 118 |
| Brown Swiss | 92 | 92 | 92 | 92 | 90 |
| Guernsey | 30 | 30 | 30 | 30 | 30 |
| N'Dama | 98 | 98 | 59 | 59 | 59 |
| Brahman | 127 | 127 | 86 | 86 | 50 |
| Nelore | 941 | 941 | 708 | 200 | 50 |
| Total | 48776 | 17813 | 17852 | 2558 | 2143 |

<sup>a</sup>Number of registered animals determined by pedigree analysis to be fullblood for breed associations with open herdbooks. <sup>b</sup>Number of registered animals assigned to their identified breed with $P \geq 0.97$ by fastSTRUCTURE in preliminary analyses and retained for subsequent analyses. <sup>c</sup>A random sample of 200 individuals was obtained for breeds with $>200$ individuals after fastSTRUCTURE analysis and all individuals were sampled for breeds with $\leq 200$ per breed and the data were again analyzed by fastSTRUCTURE with $K=19$ after removal of the Salers. <sup>d</sup>Animals that were determined to not be fullblood by pedigree analysis and animals assigned with $P \leq 0.60$ by SNPweights to their breed of registry were removed.

**Table 2.** The number of variants queried by each assay and the number of individuals from the 20 reference breeds genotyped using each assay.

| **Assay** | **No. of Variants** | **No. of Registered Individuals** |
| --- | --- | --- |
| BovineSNP50 | 58336 | 20485 |
| BovineHD | 777962 | 2303 |
| GGP-F250 | 227234 | 3068 |
| GGP-90KT | 76999 | 4407 |
| GGP-LDV3 | 26504 | 6065 |
| GGP-HDV3 | 139977 | 3630 |
| GGP-LDV4 | 30105 | 8653 |
| GGP-LDV1 | 8762 | 165 |
| Zoetis i50K | 59825 | 0 |
| ICBF IDBv3 | 53450 | 0 |
| BOVGv1 | 47843 | 0 |
| Total |  | 48776 |

### Marker set determination

To maximize the utility of the developed breed assignment tool, we identified the intersection set of SNP markers located on the bovine assays for which we had available genotype data (Table 2). However, during the process of identifying the animals that would define the breed reference panel, only 16 individuals had been genotyped using the GGP-LDV4 (n=2) and GGP-LDV3 (n=14) assays and no animals had been genotyped using the GGP-LDV1 assay. To retain as many SNP markers as possible for subsequent analysis, we identified the intersection of markers present on the GGP-90KT, GGP-F250, GGP-HDV3, GGP-LDV3, GGP-LDV4, BovineHD, BovineSNP50 and i50K assays. This intersection set included 6,799 SNP markers (BC7K). The intersection of the markers representing 5 assays (GGP-90KT, GGP-F250, GGP-HDV3, BovineHD, and BovineSNP50) was 13,291 markers (BC13K). By removing only the 16 individuals from the breed reference panel that had been genotyped on the GGP-LDV3 and GGP-LDV4 assays, we were able to compare ancestry predictions using two marker set densities (BC13K and BC7K).

### Pipeline

The developed CRUMBLER pipeline integrates the tools and the computational efficiency of publicly available software, PLINK [7,8], EIGENSOFT [9,10] and SNPweights [11] to generate ancestry estimates (Fig. 1). The pipeline integrates the often cumbersome processes of data reformatting and sequentially processing the data using analytical tools to generate ancestry proportions for targeted individuals based on a curated breed reference panel.

**Fig. 1.**
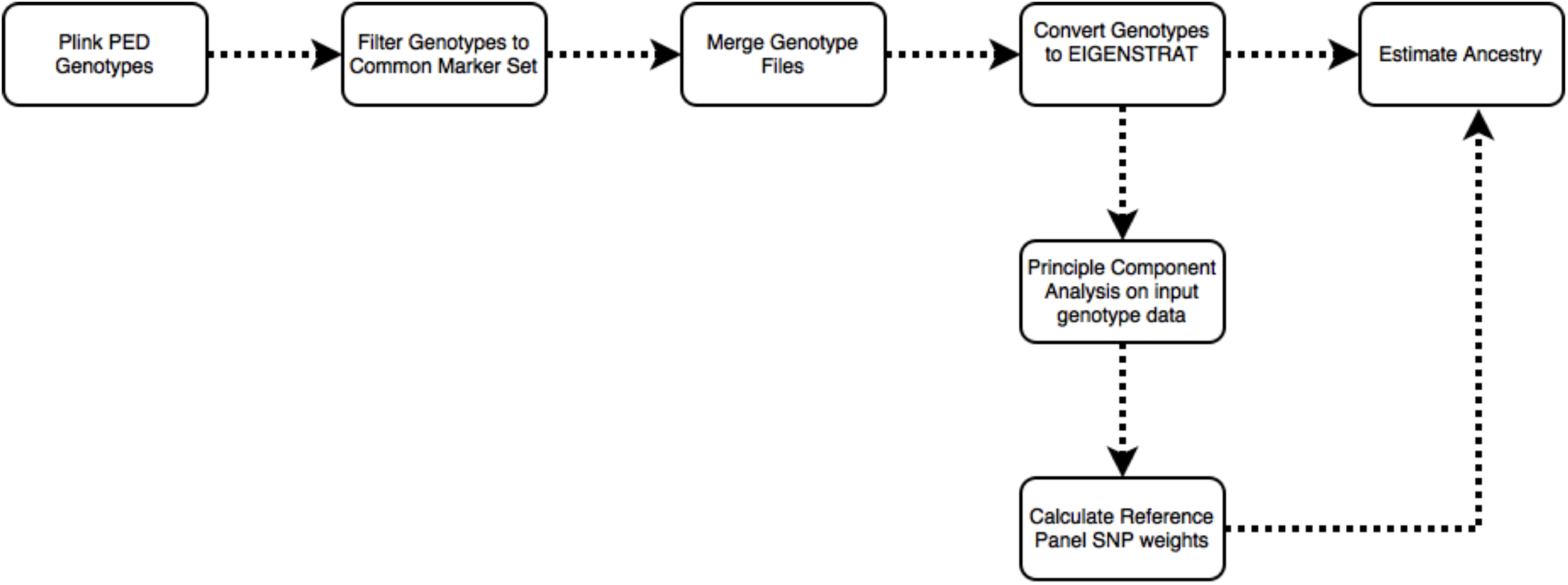
Flow diagram of the breed composition pipeline.

### PLINK

PLINK PED formatted genotypes are required as input to the pipeline. PLINK (v1.90b3.31) was used for data filtering and formatting. Genotypes can arise from any of the common bovine genotyping platforms (Table 2), provided that a PLINK compatible MAP file is provided for each assay and data produced using only a single genotyping assay is included in each PED file. The pipeline utilizes the PLINK marker filtering tool (--extract) to extract the user-specified marker subset for ancestry analysis. For analyses of animals genotyped on different genotyping platforms, the marker list representing the intersection of the platforms can be provided to extract the markers that are common to all assays. The pipeline allows multiple input genotype files and uses the PLINK merge genotype files tool (--merge) to combine genotypes into a single file for downstream analysis.

### EIGENSOFT

The EIGENSOFT convertf package is used to convert all genotypes from PLINK PED format into EIGENSTRAT format which is required by the SNPweights software. To process the reference panel data, principal component analysis using EIGENSOFT smartpca is used to generate the eigenvalues and eigenvectors that are required to calculate SNP weights using SNPweights. However, the smartpca package included in EIGENSOFT versions beyond 5.0.2 is not compatible with SNPweights. SNPweights requires an input variable, “*trace*”, to be located in the log file output from the smartpca analysis. For versions of EIGENSOFT beyond 5.0.2, the source code can be edited to ensure that the log file output is compatible with the SNPweights software (See Supplementary Information).

### SNPweights

SNPweights implements an ancestry inference model based on genome-wide SNP weights computed using genotype data for an external panel of reference individuals. To obtain SNP weights, the matrix (**g_ij_**) of reference panel genotypes for SNP **i=1, …, M** and individual **j=1, …, N** is normalized by subtracting the mean μ**_i_ =** N^-1^∑**_j_ g_ij_** and dividing by the standard deviation [**p_i_(1-p_i_)**]**^0.5^** for each SNP, where **p_i_ =** μ**_i_/2,** to improve the results of the subsequent PCA analysis from which a kinship matrix is generated [15]. A principal component decomposition is then used to generate the eigenvalues and corresponding eigenvectors of the kinship matrix [11]. The SNP weights file only needs to be recalculated if the reference panel is changed. EIGENSTRAT formatted target animal genotypes are input into SNPweights, along with the precomputed reference panel SNP weights. The SNP weights are then applied to the target individuals to estimate their ancestry proportions [11].

### Reference panel development

The definition of a set of reference individuals that define the genotype frequencies at each SNP variant for each reference breed is technically demanding, but vitally important to the process of defining ancestry. This process assumes that selection has not operated to change gene frequencies between target and reference population animals, and that each population is sufficiently large that drift has not impacted allele frequencies. It also assumes that migration between different countries does not influence population allele frequencies when registered animals are imported or exported. FastSTRUCTURE [12] analysis and iterations of animal filtering using SNPweights was performed using the genotypes of candidate reference panel individuals to remove individuals with significant evidence of admixture from the reference breed panel. An overview of the processes and iterations of filtering conducted in the development of this reference panel set is shown in Fig. S1 and Table 1.

### FastSTRUCTURE analysis to identify candidate reference panel individuals

Genotype data for 48,776 individuals produced by one of 8 different genotyping assays were available for fastSTRUCTURE analysis (Table 1) [12]. We initially performed focused fastSTRUCTURE analyses using small numbers of reference breeds including Angus and Simmental, Angus and Gelbvieh, Angus and Limousin, Angus and Red Angus, Red Angus, Hereford, Shorthorn and Salers, Red Angus, Hereford and Shorthorn, and N’Dama, Nelore and Brahman (Figs. S2-S8). Individuals possessing an ancestry assignment of at least 97% to their designated breed were retained for subsequent analysis (see Supplementary Methods and Table 1). Following filtering based on fastSTRUCTURE breed assignment, 17,852 individuals representing 19 of the original breeds remained for further analysis (Supplementary Methods and Figs. S2-S8). All of the Salers animals were removed in this filtering analysis which is consistent with previous work that found that Salers and Limousin were very similar [4]. Variation in reference population sample sizes has been shown to substantially influence the estimation of the number of ancestral populations (K) in ancestry analyses [6,13,14]. To minimize this effect and produce similar sample sizes for each of the reference breeds, we randomly sampled 200 individuals from each reference breed for which at least 200 individuals remained after filtering on an ancestry assignment of at least 97%, otherwise all remaining individuals were included for the breed (Table 1). Following fastSTRUCTURE analysis using K=19 after removal of Salers and using the BC7K marker set, Texas Longhorn was also removed from the reference panel breed list due to the inability to distinguish Texas Longhorn as a distinct population (Figure 2). Further, due to the known common ancestry [15] and similarity between Nelore and Brahman (Figure 2), the breeds were combined to represent *Bos taurus indicus*.

**Fig. 2.**
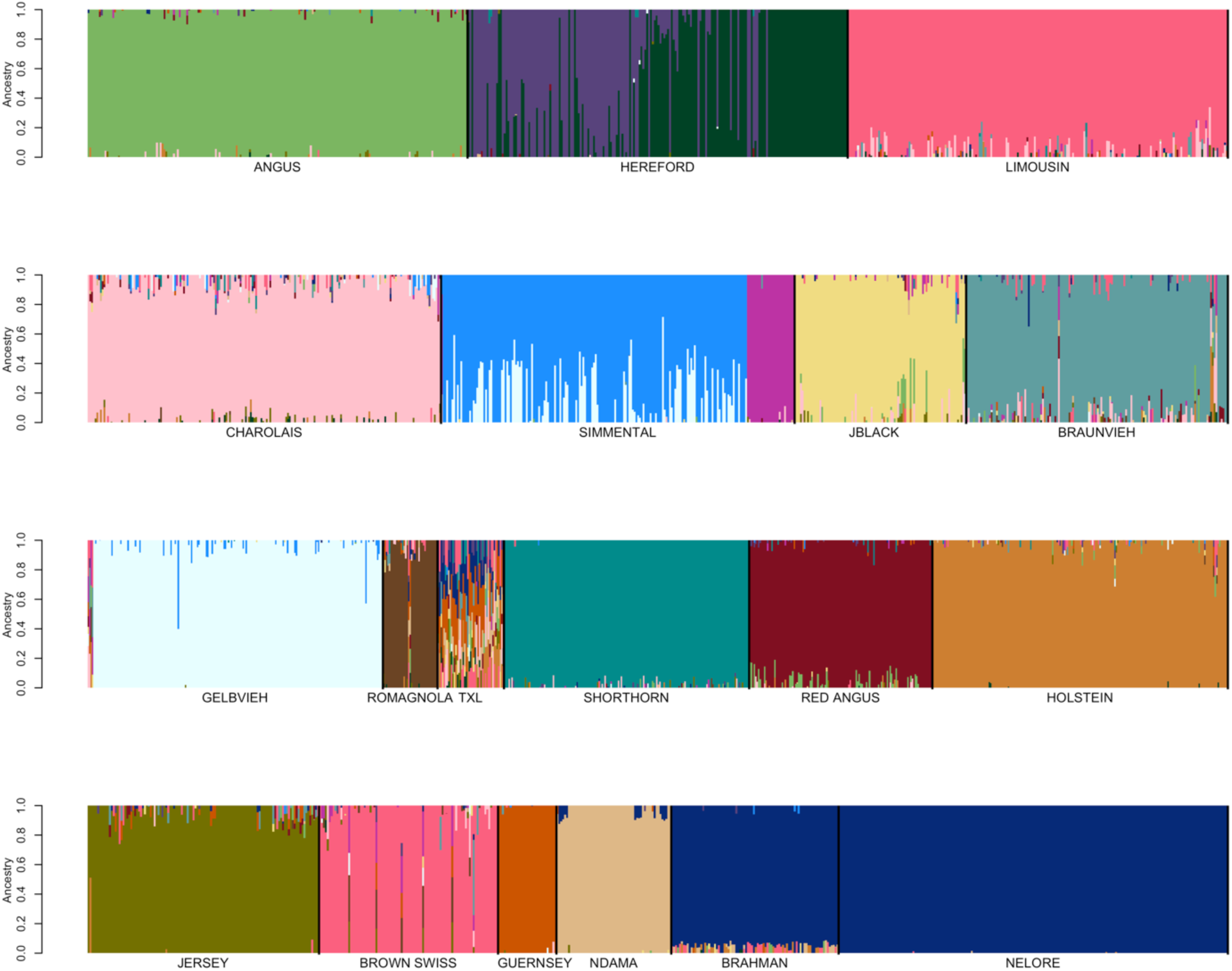
FastSTRUCTURE results for a random sample of ≤200 individuals per breed from the pool of 17,852 potential reference individuals at K=19. Breed identification is shown below each colored block and each animal is represented as a vertical line within the block.

### SNPweights analyses to refine and validate reference panel members

Random sampling of reference breed individuals was performed to create sample sets containing ≤n individuals per breed, for n = 50, 100, 150 and 200 individuals (Figs. 3 a-b and Figs. S9-S10). Sampling was performed such that if a reference breed had ≥n candidates then n individuals were randomly sampled, otherwise, all available individuals were sampled. An analysis was performed using the BC7K marker set, SNPweights was used to assign reference breed ancestries to the same sample of individuals that was used to produce the SNP weights for each of the four samples of individuals (Figs. 3 a-b and Figs. S9-S10). In the self-assignment analyses conducted using the reference breed sample sets of ≤100 individuals per breed and ≤50 individuals per breed, 7 individuals were removed due to their estimated breed ancestry being ≤60% to their registry breed (Holstein n=3, Jersey n=1, Japanese Black n=3) (Figs. 3 a-b).

**Fig. 3.**
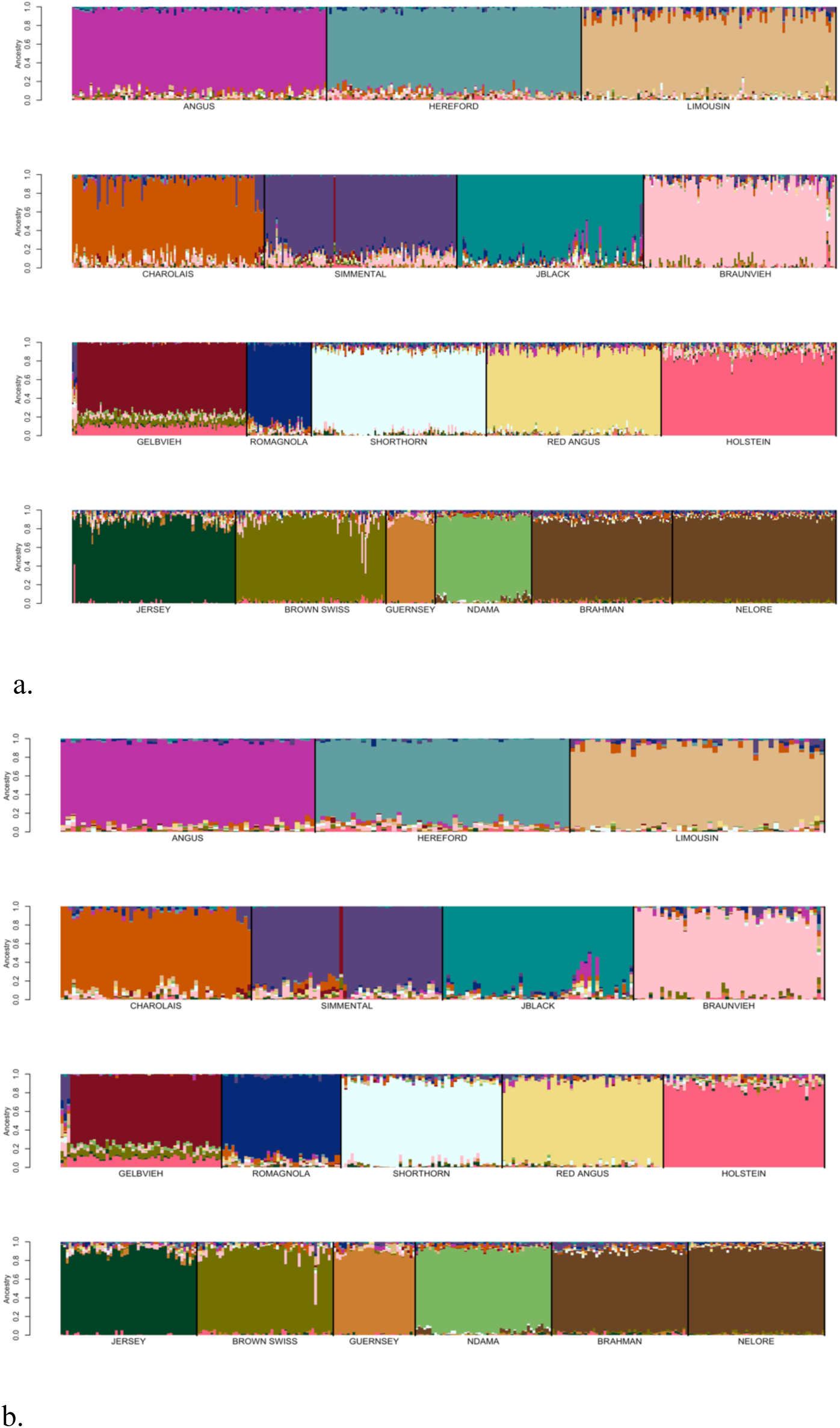
SNPweights self-assignment analysis results for reference panel sample sets consisting of: (a) ≤100 individuals per breed, or (b) ≤50 individuals per breed. Seven individuals were filtered for ≤60% ancestry to their breed of registry (Holstein n=3, Jersey n=1, Japanese Black n=3).

### Breeds with open herdbooks

For the Gelbvieh, Limousin, Shorthorn, Simmental, and Braunvieh breeds that have open U.S. herdbook registries, fullblood or 100% ancestry individuals were identified based on pedigree data obtained from the respective breed associations (Table 1). The term “fullblood” is used to identify cattle for which every ancestor is registered in the herdbook and can be traced back to the breed founders. The term “purebred” refers to animals that have been graded up via crossbreeding to purebred status. Charolais also has an open herdbook registry in the U.S., however, access to Full French imported Charolais breed members was limited. As a result, all Charolais individuals identified as purebred in the association registry were retained for downstream analysis, however, these individuals could contain up to 1/32 introgression from another breed. A random sample of 200 individuals was taken for each breed with more than 200 identified fullblood individuals, otherwise all animals were sampled. Individuals previously included in the candidate reference panel following preliminary fastSTRUCTURE filtering for the open herd book breeds were removed and replaced with the fullblood individuals.

### Additional reference panel filtering using SNPweights

After filtering animals identified to not be fullblood based on their pedigree information, we randomly sampled ≤50 individuals per reference breed and utilized SNPweights to estimate weights for each sample and also to estimate breed ancestries for members of the same sample that was used to generate the SNP weights. Based on these analyses, we created 5 overlapping reference breed sets, each containing individuals with ≥90%, ≥85%, ≥80%, ≥75%, or ≥70% ancestry assignment to their registry breeds (Table 3).

**Table 3.** Number of individuals for each reference breed assigned to their breed of registration by minimum ancestry threshold.

| Breed | Breed Assignment Probability |  |  |  |  |
| --- | --- | --- | --- | --- | --- |
|  | ≥90% | ≥85% | ≥80% | ≥75% | ≥70% |
| Angus | 51 | 136 | 184 | 199 | 200 |
| Hereford | 58 | 136 | 184 | 200 | 200 |
| Limousin | 93 | 127 | 144 | 162 | 173 |
| Charolais | 52 | 92 | 119 | 132 | 147 |
| Simmental | 21 | 43 | 81 | 103 | 121 |
| Japanese Black | 52 | 73 | 78 | 83 | 86 |
| Braunvieh | 37 | 57 | 63 | 65 | 68 |
| Gelbvieh | 23 | 31 | 39 | 43 | 43 |
| Romagnola | 10 | 25 | 32 | 36 | 37 |
| Shorthorn | 34 | 98 | 159 | 170 | 177 |
| Red Angus | 48 | 88 | 110 | 120 | 123 |
| Holstein | 39 | 119 | 172 | 193 | 196 |
| Jersey | 52 | 77 | 91 | 108 | 116 |
| Brown Swiss | 38 | 64 | 73 | 82 | 86 |
| Guernsey | 12 | 22 | 29 | 30 | 30 |
| N'Dama | 27 | 45 | 59 | 59 | 59 |
| Brahman | 15 | 40 | 50 | 50 | 50 |
| Nelore | 32 | 50 | 50 | 50 | 50 |
| Total | 694 | 1323 | 1717 | 1885 | 1962 |

### Simulated Genotypes

Using the phased BC7K genotypes for the final reference population of 803 individuals (3 Nelore genotyped with the BovineHD assay were removed because they were determined to cause problems for the phasing software), we simulated genotypes for 803 individuals each generation (N = 1, 3, 5 and 10) by randomly sampling two individuals as parents from generation N-1 and using a Poisson distribution to sample at random a single recombinant chromosome from each parent. The number of recombination events for each sampled chromosome was sampled from a Poisson distribution with mean equal to chromosome length in Mb/100 (i.e. 1.58 Morgans for chromosome 1). Simulated genotypes were produced for individuals 1 generation removed from the fullblood/purebred reference population animals (i.e., 50% breed A and 50% breed B), 3, 5, and 10, generations, respectively, to evaluate the ability of CRUMBLER to detect large through to small admixture proportions in animals with increasing numbers of breeds represented in their ancestry. Breed composition estimates for these animals were obtained by tracing the breed of origin of every allele present in each generation N animal. For each marker, we attributed the genomic fragment from the center points of the intervals on each side of each marker to the breed of origin of the two alleles at each marker and summed these across all loci. Finally, we normalized these sums by dividing by the autosomal genome size using UMD3.1 coordinates.

## Results and Discussion

The concept of breed and breed membership is man-made and does not inherently exist in nature. Moreover, the formation of breeds of cattle is very recent, as cattle domestication began about 10,000 years ago but the formation of herdbooks has occurred only during the last 200-250 years [16]. Nevertheless, the effects of drift and human selection over the last 200 years have caused sufficient divergence among breeds that breed differences are identifiable at the molecular level. Such signals are essential for breed ancestry analyses to be effective in modern admixed animals. Previous work on assigning breed composition in admixed cattle utilized 50K genotype data and a reference panel of 16 breeds, with the basis for reference panel inclusion being breed association registration [17]. However, the continual evolution of genotyping assays has led to content changes resulting in only a relatively small proportion of markers in common among assays. Consequently, there is a need to evaluate whether these markers are sufficient for breed content estimation, leading to their conservation in the design of future assays. Furthermore the development of an analytical pipeline based on these markers would simplify analysis for end-users and the use of a single reference panel would allow the direct comparison of results between applications.

### Reference panel development

Previously developed cattle reference panels have relied on pedigree accuracy and breed association registration for their definition [17]. Conversely, we used an iterative approach for reference population curation that was able to validate the accuracy of the pedigree information used to identify candidates. FastSTRUCTURE analyses performed using the candidate individuals for each of the initial 19 reference breeds suggested population subdivision in both the Hereford and Simmental (Figure 2). Pedigree analysis for the Herefords within each subpopulation indicated that the subpopulations comprised animals from the highly inbred USDA Miles City Line 1 Hereford population (L1) and other individuals representing broader U.S. Hereford pedigrees. The Miles City L1 Hereford cattle were derived from two bulls, both sired by Advance Domino 13 (AHA registration number 1668403) and 50 Hereford foundation cows. Since the founding of the L1 Herefords, the migration of germplasm has been unidirectional from L1 into the broader U.S. industry, as the L1 population has been closed since its founding [18]. However, the L1 Herefords have profoundly influenced the U.S. Hereford population. L1 Herefords do not segregate for recessive dwarfism, which has been a threat to Hereford breeders since the 1950s, and this has led to L1 cattle becoming popular in the process of purging herds of the defect [19]. In 1980, the average proportion of U.S. registered Herefords influenced by L1 genetics was 23%. By 2008, this proportion had increased to 81% [18].

The detected subpopulation division within the Simmental breed (Figure 2) represents the differentiation between purebred and fullblood animals. For example, progeny of a popular fullblood Simmental sire are present in both subpopulations, however, in one subpopulation the family members are all fullblood and in the other they are all purebred or percentage Simmental animals. This result supports the need to identify fullblood animals as reference panel breed representatives for breeds with open herdbooks.

#### Reference population sample size

By randomly sampling individuals from the candidate reference breed set and using SNPweights to assign these individuals to reference populations, we found that reference panel breed sample sizes of ≤50 or ≤100 individuals appeared to capture the diversity within each breed and appropriately determined the ancestry of the tested individuals (Fig. 3 a-b). For each breed, the percent ancestry predicted for the tested reference samples was, on average, 3.86% higher when the SNP weights were estimated using ≤50 individuals per breed than when ≤100 individuals per breed were used (Table 4). This reflects the increased homogeneity of individuals within each breed and a greater genetic distance between individuals from different breeds as smaller samples of individuals from each breed are used to define the reference panel. Further, due to limitations in the number of genotyped individuals for some breeds (Table 1), as the sample size was increased globally, imbalances were created between the reference panel breed sample sizes which impacted breed composition estimation (Fig. S9-S10). It has previously been shown that the power to detect population structure improves as the reference population sample sizes become more similar [6,14].

**Table 4.** Ancestry proportion statistics for the self-assignment of reference panel members from samples of ≤50 or ≤100 individuals from the candidate reference breed individuals.

| **Breed** | **Min %**<br>**($\leq 50$)** | **Avg %**<br>**($\leq 50$)** | **Max %**<br>**($\leq 50$)** | **Min %**<br>**($\leq 100$)** | **Avg %**<br>**($\leq 100$)** | **Max %**<br>**($\leq 100$)** |
| --- | --- | --- | --- | --- | --- | --- |
| Angus | 86.22 | 90.40 | 95.54 | 78.49 | 87.05 | 94.13 |
| Hereford | 79.75 | 90.08 | 95.05 | 73.41 | 87.39 | 96.81 |
| Limousin | 69.52 | 88.53 | 98.16 | 18.36 | 86.40 | 98.81 |
| Charolais | 78.14 | 90.19 | 99.82 | 48.93 | 77.46 | 93.96 |
| Simmental | 81.06 | 90.37 | 97.66 | 61.36 | 73.05 | 88.11 |
| Japanese Black | 81.44 | 90.00 | 97.07 | 24.51 | 86.50 | 98.95 |
| Braunvieh | 71.59 | 89.46 | 98.61 | 65.46 | 88.36 | 98.70 |
| Gelbvieh | 73.03 | 76.27 | 81.63 | 60.92 | 74.59 | 80.33 |
| Romagnola | 75.05 | 87.18 | 96.66 | 74.79 | 85.99 | 95.12 |
| Shorthorn | 84.42 | 88.69 | 94.54 | 70.71 | 85.27 | 96.35 |
| Red Angus | 79.00 | 89.60 | 96.33 | 68.07 | 86.83 | 97.38 |
| Holstein | 85.82 | 90.30 | 97.51 | 62.95 | 86.97 | 97.81 |
| Jersey | 78.55 | 89.28 | 95.93 | 61.23 | 86.54 | 97.18 |
| Brown Swiss | 80.10 | 89.22 | 96.40 | 61.68 | 86.02 | 98.42 |
| Guernsey | 79.53 | 89.19 | 95.85 | 77.40 | 88.31 | 94.36 |
| N'Dama | 80.67 | 89.25 | 96.90 | 78.91 | 87.78 | 95.67 |
| *Bos taurus indicus* | 87.83 | 91.91 | 97.75 | 81.43 | 89.79 | 97.60 |

#### Marker density

After the replacement of reference breed individuals with those identified to be fullblood based on pedigree analysis for the open herdbook Gelbvieh, Simmental, Limousin, Braunvieh, Shorthorn, and Charolais breeds, additional self-assignment analyses were conducted to evaluate the effects of marker set size on ancestry prediction. Breed reference panels were again constructed by randomly sampling ≤50 individuals per breed and SNP weights were calculated using both the BC13K markers and BC7K markers. The estimated SNP weights were then used to self-assign ancestry to members of the reference panel animals representing the reference breed set. The ancestry predictions for the reference breed individuals using either the BC7K (Fig. 4a; Fig. S11) or BC13K (Fig. 4b; Fig. S12) marker sets indicate that use of the BC13K marker set did not significantly impact the ancestry predictions. Consequently, the use of the 6,799 markers common to the 8 commercially available genotyping platforms appears to be sufficient to assign breed ancestry for the majority of animals produced in the U.S. The CRUMBLER pipeline can accommodate samples genotyped using alternative assays, however, the produced breed composition estimates will be based on the intersection of markers on the assay and the BC7K marker set.

**Fig. 4.**
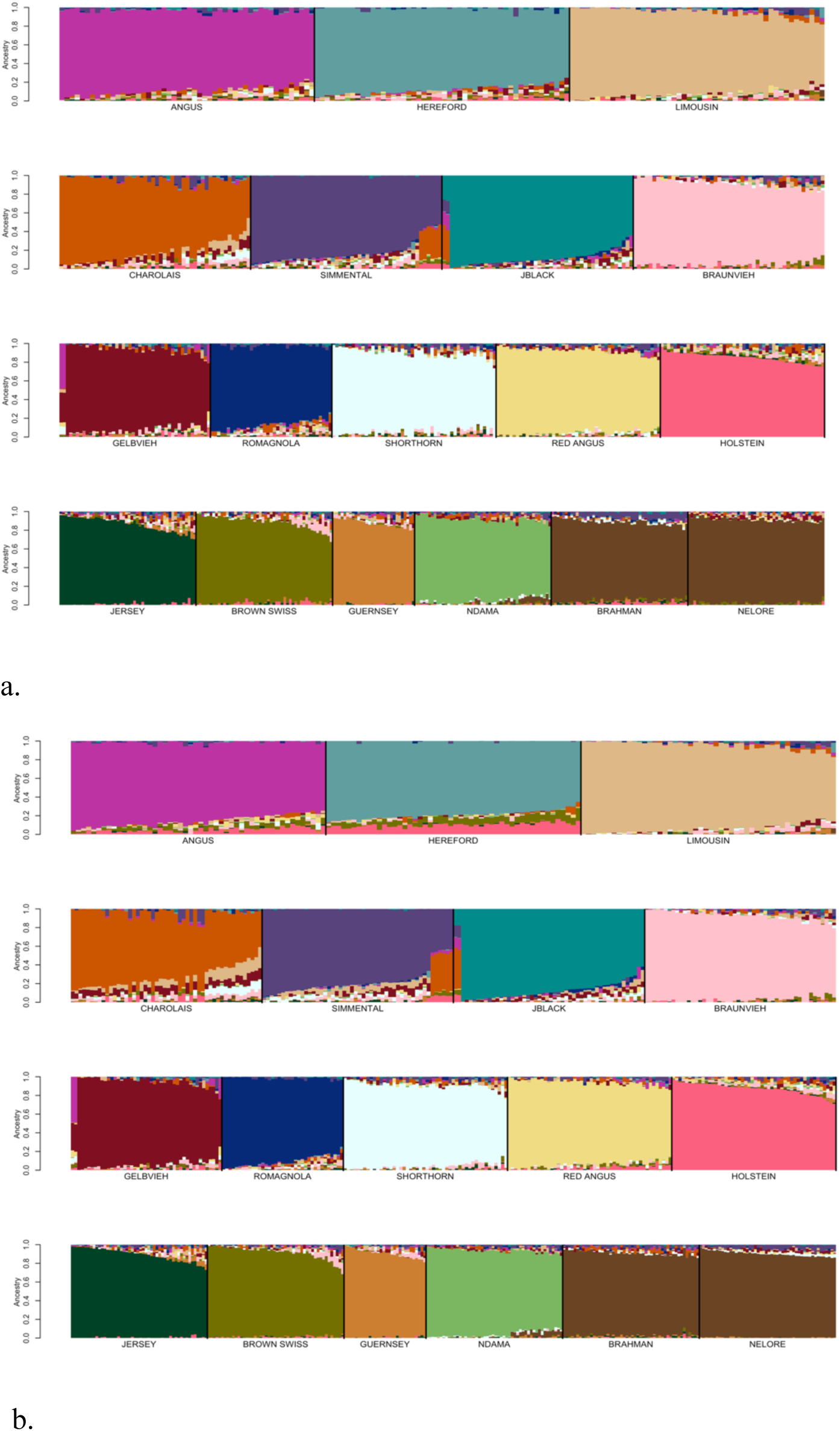
SNPweights self-assignment of ancestry for candidate reference breed individuals following evaluation of open herdbook breeds using: (a) the BC7K, or (b) the BC13K marker panels. Reference breed panels were constructed by random sampling ≤50 individuals per breed and SNP weights were estimated using the BC7K and BC13K marker sets.

#### Assignment thresholds

We next examined the effects of reference breed homogeneity on ancestry assignment by identifying reference panel members that had been assigned to their breed of registry using SNPweights with probabilities of ancestry of ≥90%, ≥85%, ≥80%, ≥75%, and ≥70%, respectively (Table 3). From these individuals, reference breed panels were obtained by randomly sampling ≤50 individuals per breed, until each individual was represented in at least one sample set. SNP weights were then estimated using the BC7K marker set and ancestry was assigned for these individuals using SNPweights (Figs. 5-6 and Figs. S13-S15). Limiting the reference breed panel members to those individuals with ≥90% ancestry assigned to their breed of registry produced a reference panel that did not represent the extent of diversity within each of the breeds (Fig. 5). On the other hand, using an ancestry assignment of ≥85% clearly captured greater diversity within each breed (Fig. 6) and maximized the self-assignment of ancestry to the breed of registration (Table 5).

**Fig. 5.**
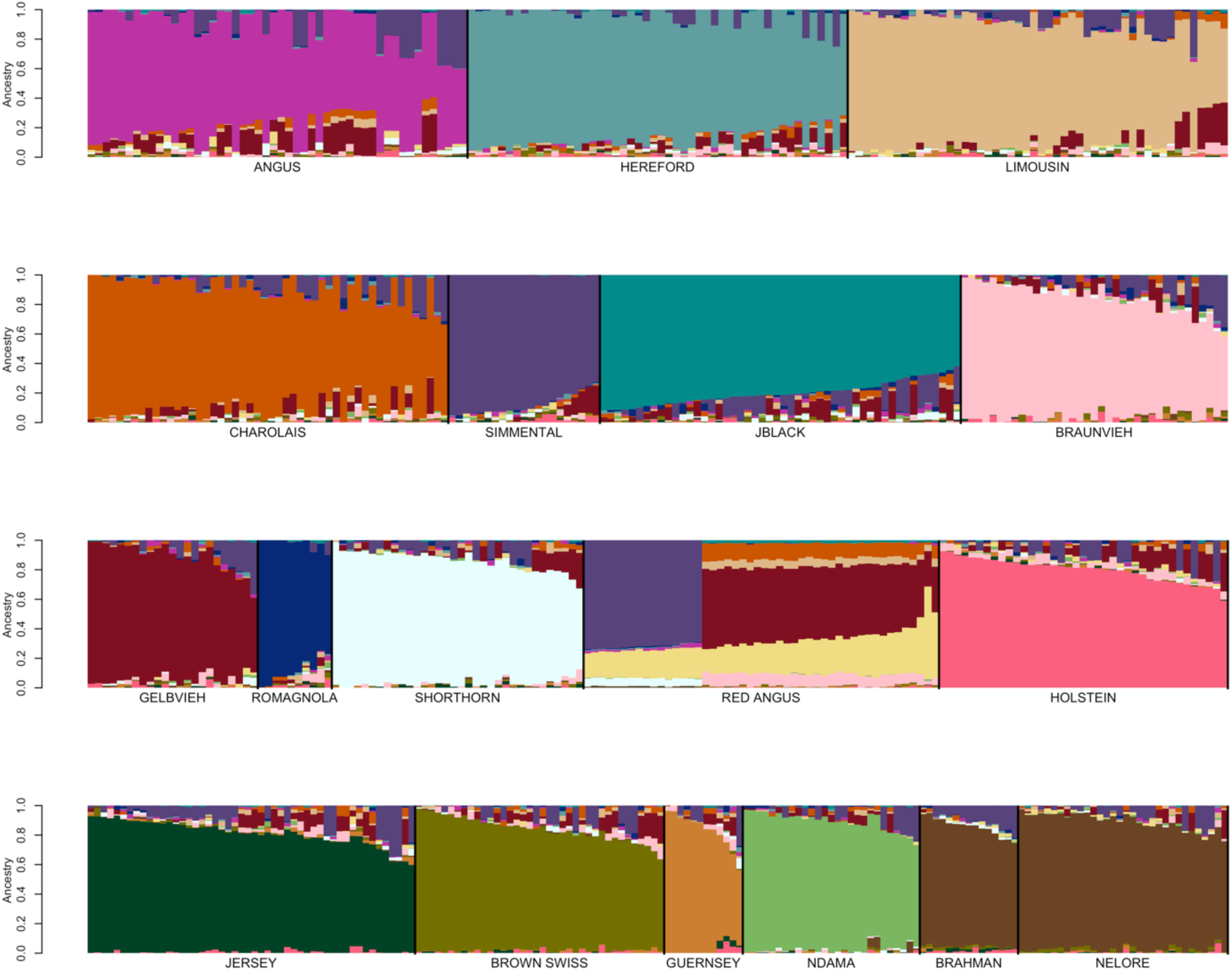
Reference breed panel constructed by the random sampling of ≤50 individuals per breed from individuals with ≥90% ancestry was self-assigned to reference breed ancestry using the BC7K marker set.

**Fig. 6.**
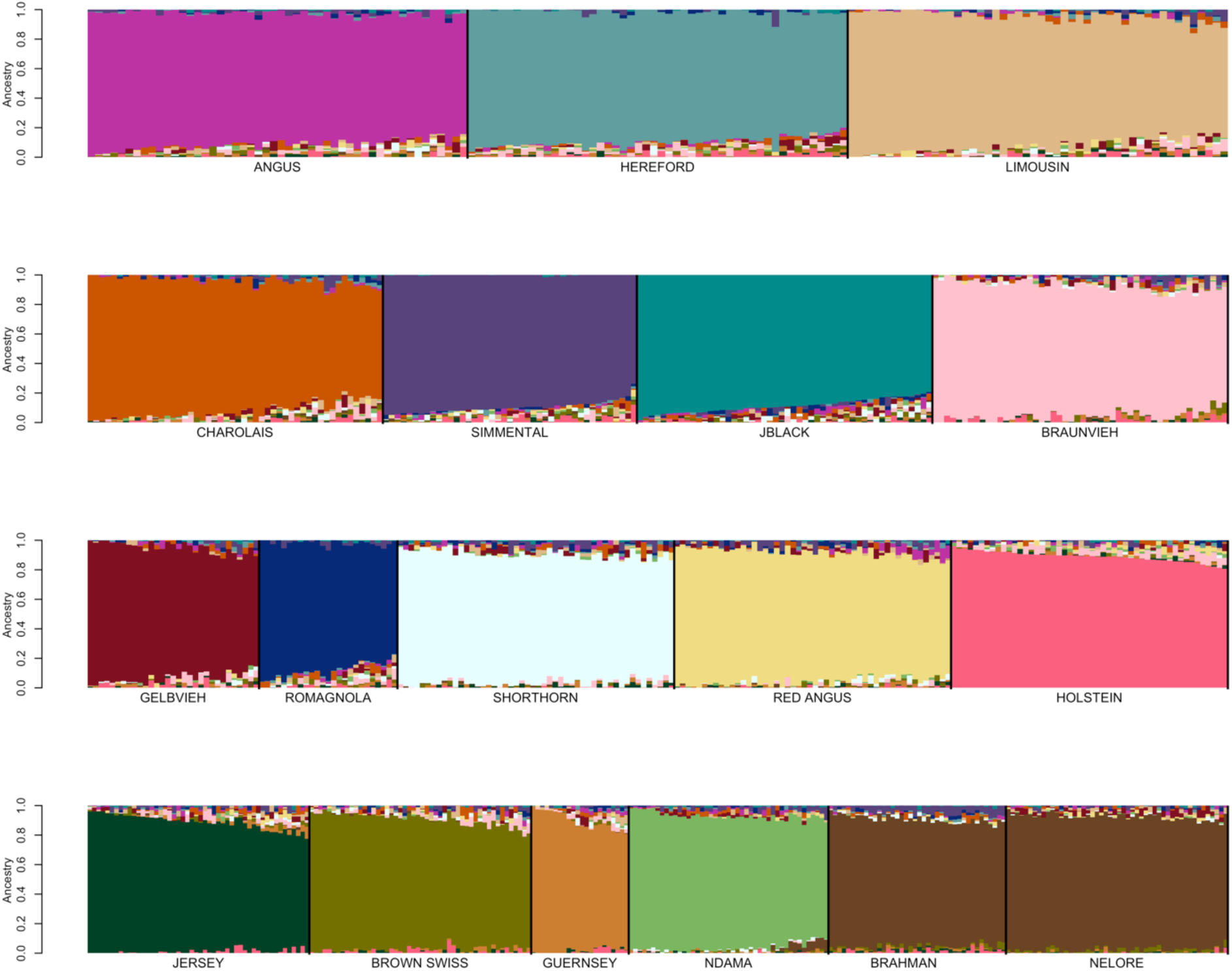
Reference breed panel constructed by the random sampling of ≤50 individuals per breed from individuals with ≥85% ancestry was self-assigned to reference breed ancestry using the BC7K marker set.

**Table 5.**
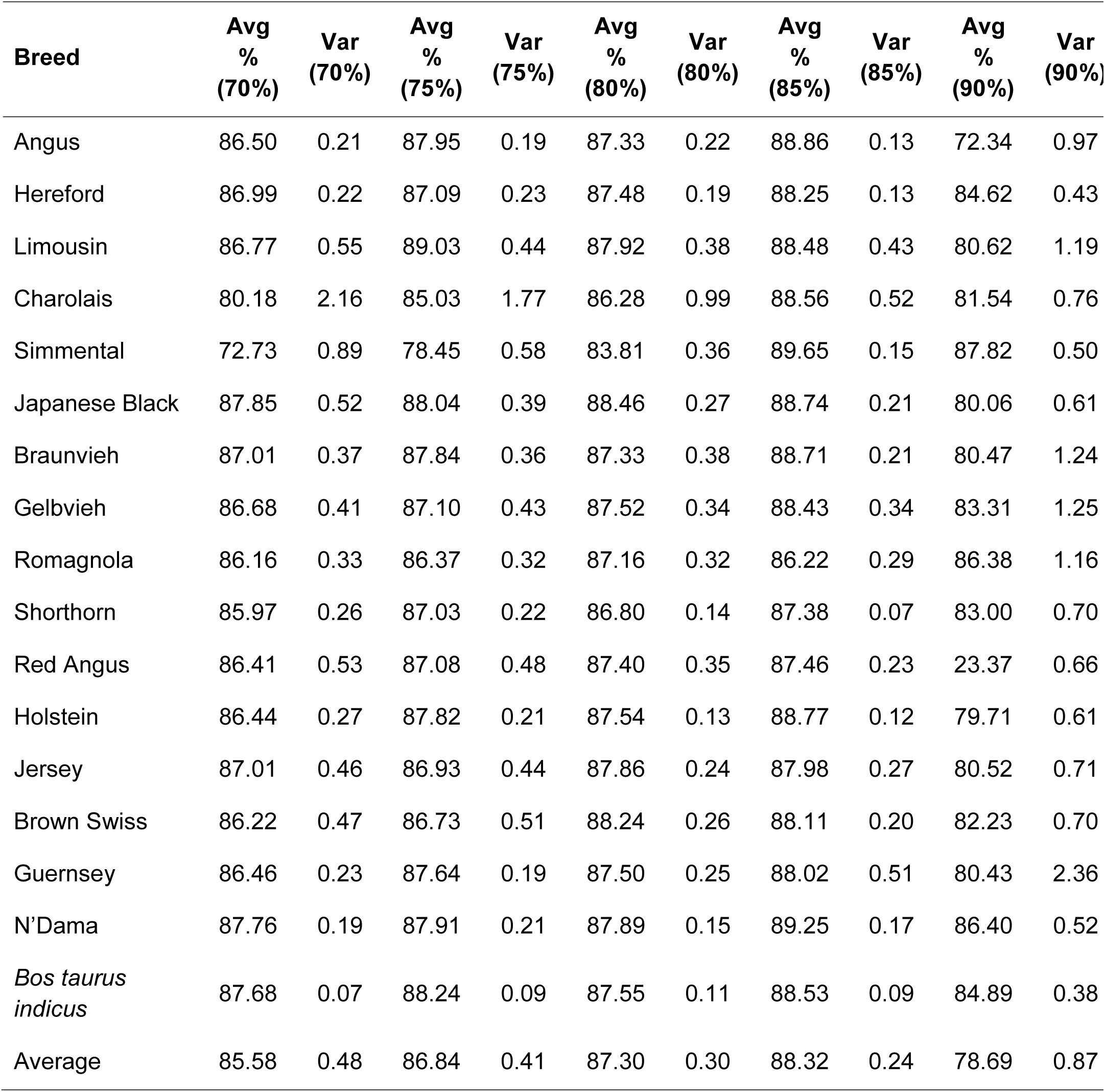
Average predicted ancestry and variance in predicted ancestry for candidate reference breed individuals when filtered on minimum predicted ancestry.

| Breed | Avg<br>%<br>(70%) | Var<br>(70%) | Avg<br>%<br>(75%) | Var<br>(75%) | Avg<br>%<br>(80%) | Var<br>(80%) | Avg<br>%<br>(85%) | Var<br>(85%) | Avg<br>%<br>(90%) | Var<br>(90%) |
| --- | --- | --- | --- | --- | --- | --- | --- | --- | --- | --- |
| Angus | 86.50 | 0.21 | 87.95 | 0.19 | 87.33 | 0.22 | 88.86 | 0.13 | 72.34 | 0.97 |
| Hereford | 86.99 | 0.22 | 87.09 | 0.23 | 87.48 | 0.19 | 88.25 | 0.13 | 84.62 | 0.43 |
| Limousin | 86.77 | 0.55 | 89.03 | 0.44 | 87.92 | 0.38 | 88.48 | 0.43 | 80.62 | 1.19 |
| Charolais | 80.18 | 2.16 | 85.03 | 1.77 | 86.28 | 0.99 | 88.56 | 0.52 | 81.54 | 0.76 |
| Simmental | 72.73 | 0.89 | 78.45 | 0.58 | 83.81 | 0.36 | 89.65 | 0.15 | 87.82 | 0.50 |
| Japanese Black | 87.85 | 0.52 | 88.04 | 0.39 | 88.46 | 0.27 | 88.74 | 0.21 | 80.06 | 0.61 |
| Braunvieh | 87.01 | 0.37 | 87.84 | 0.36 | 87.33 | 0.38 | 88.71 | 0.21 | 80.47 | 1.24 |
| Gelbvieh | 86.68 | 0.41 | 87.10 | 0.43 | 87.52 | 0.34 | 88.43 | 0.34 | 83.31 | 1.25 |
| Romagnola | 86.16 | 0.33 | 86.37 | 0.32 | 87.16 | 0.32 | 86.22 | 0.29 | 86.38 | 1.16 |
| Shorthorn | 85.97 | 0.26 | 87.03 | 0.22 | 86.80 | 0.14 | 87.38 | 0.07 | 83.00 | 0.70 |
| Red Angus | 86.41 | 0.53 | 87.08 | 0.48 | 87.40 | 0.35 | 87.46 | 0.23 | 23.37 | 0.66 |
| Holstein | 86.44 | 0.27 | 87.82 | 0.21 | 87.54 | 0.13 | 88.77 | 0.12 | 79.71 | 0.61 |
| Jersey | 87.01 | 0.46 | 86.93 | 0.44 | 87.86 | 0.24 | 87.98 | 0.27 | 80.52 | 0.71 |
| Brown Swiss | 86.22 | 0.47 | 86.73 | 0.51 | 88.24 | 0.26 | 88.11 | 0.20 | 82.23 | 0.70 |
| Guernsey | 86.46 | 0.23 | 87.64 | 0.19 | 87.50 | 0.25 | 88.02 | 0.51 | 80.43 | 2.36 |
| N'Dama | 87.76 | 0.19 | 87.91 | 0.21 | 87.89 | 0.15 | 89.25 | 0.17 | 86.40 | 0.52 |
| *Bos taurus indicus* | 87.68 | 0.07 | 88.24 | 0.09 | 87.55 | 0.11 | 88.53 | 0.09 | 84.89 | 0.38 |
| Average | 85.58 | 0.48 | 86.84 | 0.41 | 87.30 | 0.30 | 88.32 | 0.24 | 78.69 | 0.87 |

#### Reference panel definition

To examine whether the specific individuals represented in the reference panel sample influenced the self-assignment of ancestry to the sampled individuals, a second sample of ≤50 distinct individuals per breed was obtained from the individuals with ≥85% assignment to their breed of registration and analyzed with SNPweights (Fig. 7). Fig. 7 indicates that the ability to predict ancestry was not influenced by the specific individuals sampled from the set of animals with ≥85% ancestry to their breed of registration.

**Fig. 7.**
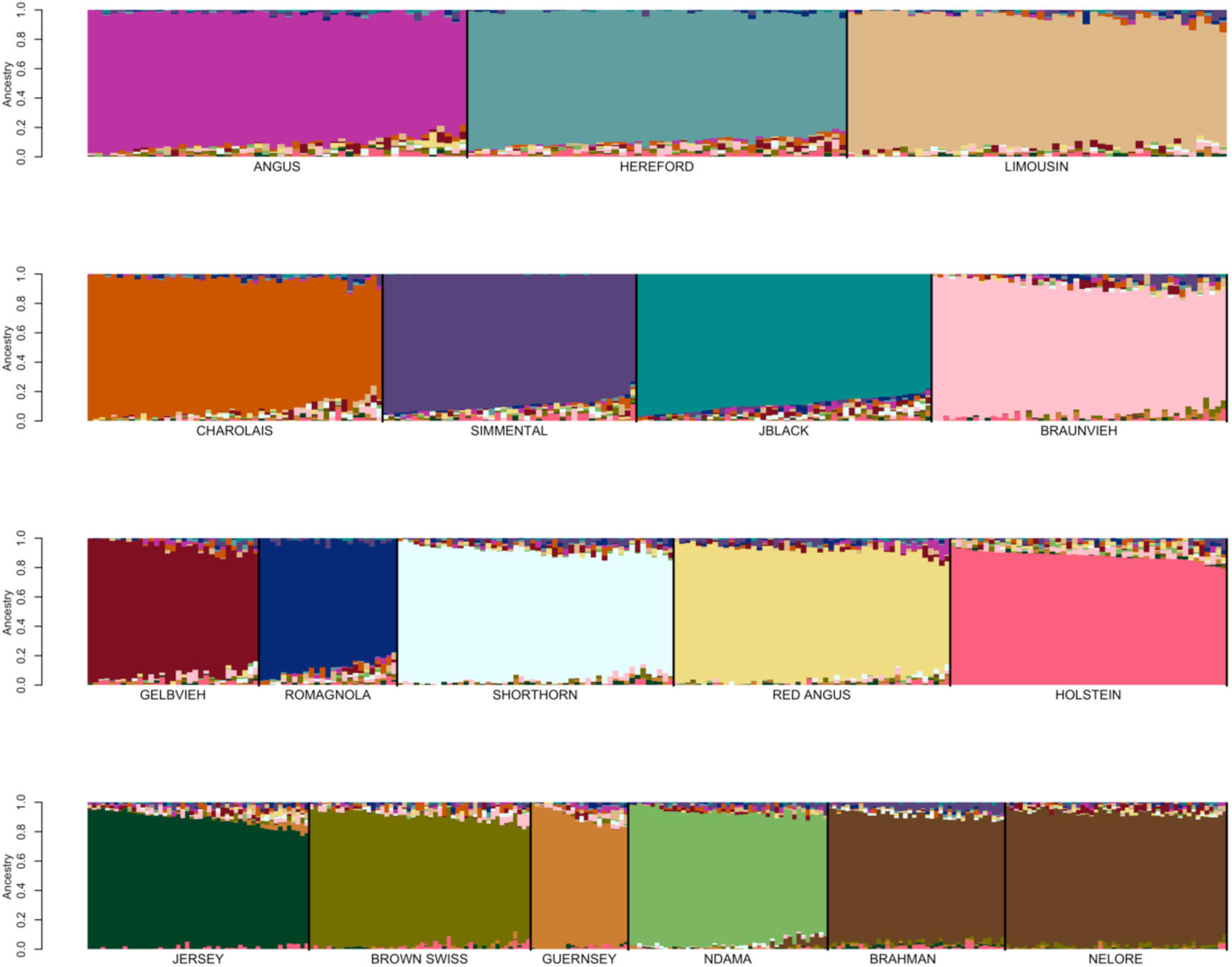
Reference breed panel constructed by the independent random sampling of a second sample of ≤50 individuals per breed from individuals with ≥85% ancestry after eliminating individuals represented in the first sample was self-assigned to reference breed ancestry using the BC7K marker set.

Additionally, Figs. 6 and 7 suggest that the use of a reference breed panel constructed by the random sampling of ≤50 individuals per breed from individuals with ≥85% self-assigned ancestry to their breed of registration maintained sufficient within-breed diversity to accurately estimate the ancestry of target individuals. However, these figures also reveal small amounts of apparent introgression from other reference panel breeds within each of the breeds. This does not appear to be an issue of marker resolution since the analyses performed with the BC7K and BC13K marker sets generated similar results (Fig. 4). We conclude that these apparent introgressions are either due to a lack of power to discriminate among breeds using the common markers designed onto commercial genotyping platforms, or represent the presence of common ancestry among the breeds prior to the formation of breed herdbooks ~200 years ago. Molecular evidence for this shared ancestry exists, for example, Hereford and Angus cattle share the *Celtic* polled allele [20] and the segmental duplication responsible for the white anterior, ventral and dorsal coat color pattern occurs only in Hereford and Simmental cattle and their crosses [21]. These data clearly indicate that crossbreeding was widespread prior to the formal conceptualization of breeds.

### Reference panel validation

To evaluate the ability of the selected reference breed panel to identify breed composition, an analysis was conducted for all 170,544 samples in the database which required 60 processor minutes (Fig 6-7). We extracted animals with pedigree information including fullblood and purebred animals registered with open herdbook breed associations and 2,243 crossbred animals with varying degrees of admixture. Considering the amount of available data, the number of pedigreed admixed animals was very limited and the purebred animals all had similar expected admixture proportions. Consequently, we next simulated genotypes for animals by assuming the random mating of members of the reference breed panel for 1, 3, 5 and 10 generations assuming non-overlapping generations to generate generations of animals with different numbers of breeds and breed proportions represented in their genomes.

#### Registered fullblood animals

For the Gelbvieh, Limousin, Shorthorn, Simmental, and Braunvieh breeds that have open herdbook registries, fullblood or 100% ancestry individuals were identified based on pedigree data obtained from the respective breed associations (Table 1). CRUMBLER estimates were obtained for these fullblood individuals and the distribution of estimates by breed are in Fig. 8. For all breeds except Charolais, >50% of the individuals had CRUMBLER estimated percentages of ≥80% to their respective breeds.

**Fig. 8.**
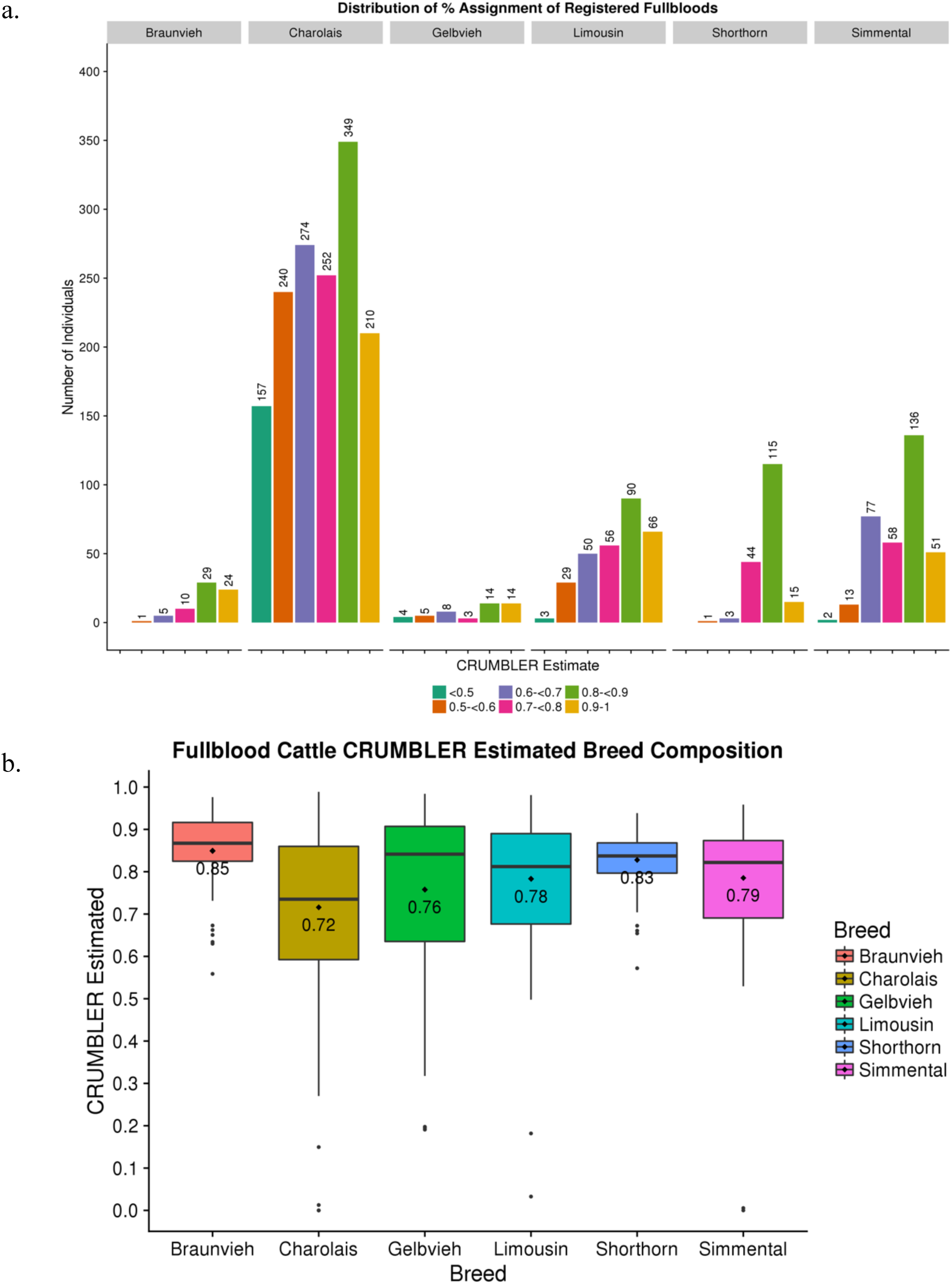
(a) Distribution by breed of SNPweights ancestry assignment results for 2,408 registered fullblood animals from open herd book breeds. (b) Pictorial representation of CRUMBLER estimates for 2,408 registered fullblood animals from open herd book breeds.

Average percentage estimates for fullblood Gelbvieh, Limousin, Shorthorn, Simmental, and Braunvieh individuals were 76%, 78%, 83%, 79%, and 85%, respectively (Fig. 8b). However, the number of genotyped imported Full French Charolais animals was limited and so we also analyzed all purebred Charolais individuals which could contain up to 1/32^nd^ of their genome introgressed from another breed. The average Charolais breed assignment was 72% and the distribution of estimates was more variable than for the fullblood animals from the other breeds (Fig. 8b).

#### Pedigreed crossbred animals

Based on pedigree, 2,005 individuals were identified as being primarily Hereford but with varying degrees of Red Angus, Salers, Angus or unknown other breed influence. The analysis results agreed with the pedigree data (Fig. 9a) To investigate the correlations between pedigree and CRUMBLER estimated breed proportions, we removed proportions for breeds that were less than 3% and normalized the remaining values. CRUMBLER estimates were then correlated with the pedigree predicted estimates of the proportion of Hereford in these individuals (Fig. 9c). CRUMBLER tended to underestimate the Hereford proportion as the pedigree estimated Hereford proportion tended to 100%.

**Fig. 9.**
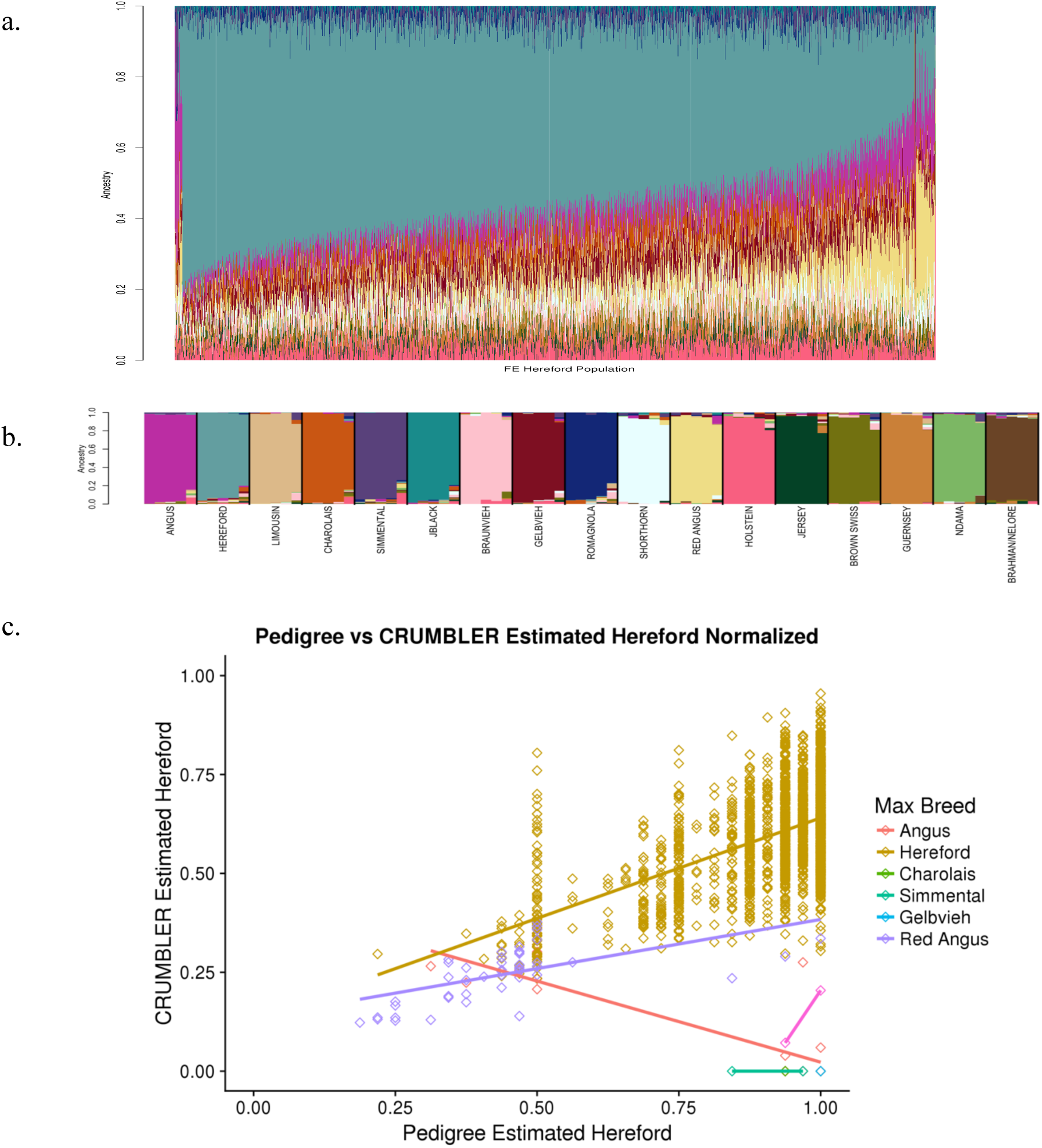
(a) SNPweights ancestry results for 2,005 crossbred Hereford individuals with *a-priori* breed composition estimates determined by pedigree. (b) Breed assignment reference breed key. (c) Hereford SNPweights estimated proportions using CRUMBLER are plotted against the pedigree estimates. Data point color indicates the breed for which SNPweights assigned the highest proportion for each individual.

The remaining 238 crossbred individuals were commercial, advanced generation animals with an expected 50% Angus and 50% Simmental ancestry based on pedigree data. Results of the CRUMBLER analysis again support the pedigree data (Fig. 10). The presence of Red Angus ancestry in these animals reveals the inability of the analysis to fully differentiate between Angus and Red Angus, which only diverged in the U.S. in 1954, and also the influence of Red Angus in the U.S. Simmental breed (Fig. S16).

**Fig. 10.**
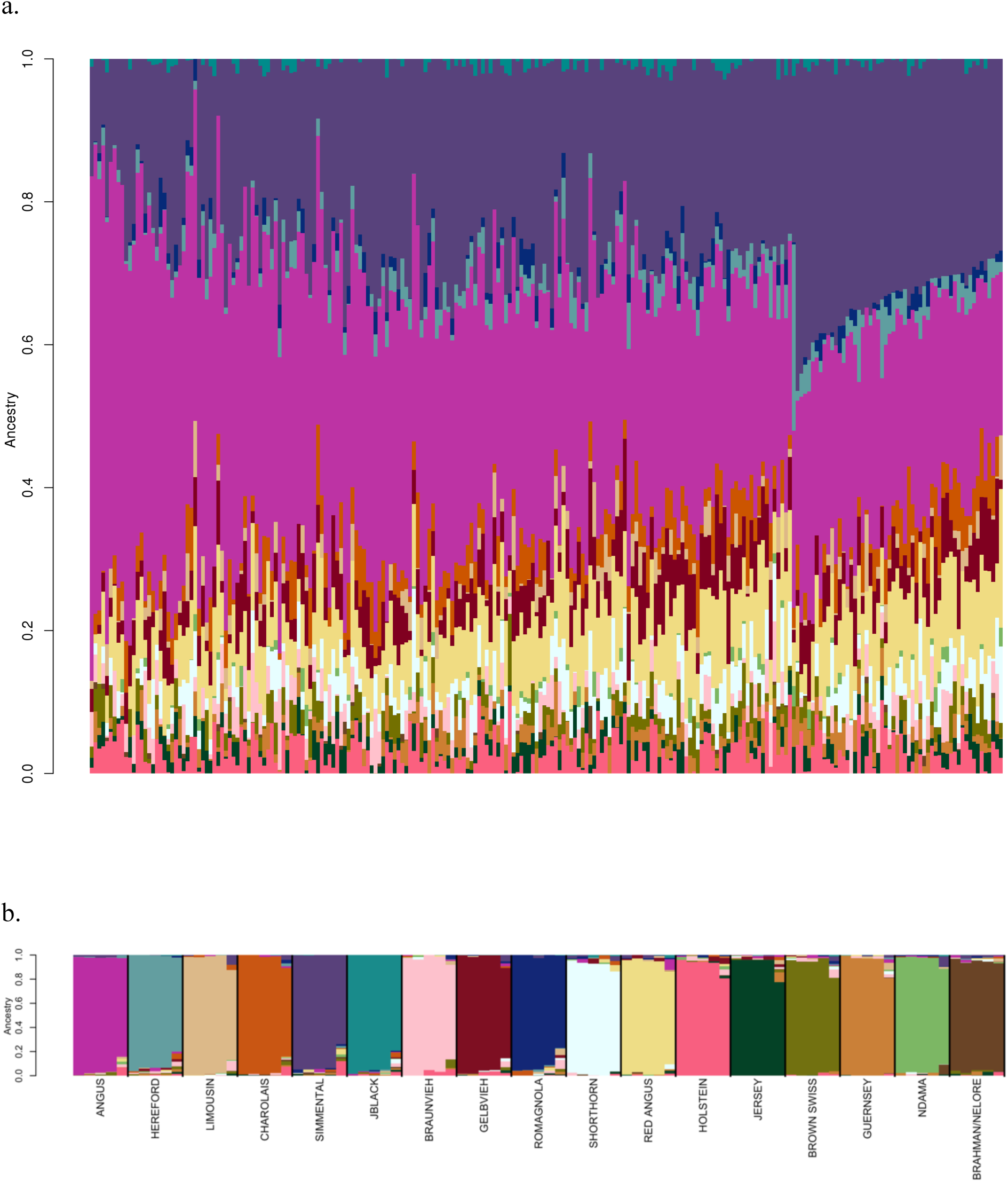
(a) SNPweights ancestry results for 238 crossbred individuals with *a-priori* breed composition estimates of 50% Angus and 50% Simmental based on a reference panel with ≤50 individuals per breed sampled from individuals with ≥85% assignment to their breed of registry. (b) Breed assignment for the crossbred individuals can be determined using this reference breed key.

#### Simulated genotypes

Genomes were simulated using the phased genotypes for 803 individuals from the reference breed panel to contain varying breed numbers and admixture proportions after 1, 3, 5, and 10 generations of random mating with nonoverlapping generations. In generation 1, the admixed individuals were F_1_ individuals with a 50:50 autosomal genome composition unless both parents were randomly sampled from the same breed. CRUMBLER estimates of breed composition using the simulated genotypes were strongly correlated with the simulated compositions, especially for generations 1 and 3 (Fig 11). As the number of generations increased, the number of breeds represented in the simulated genomes tended to increase and the proportion of the genome originating from any one breed tended to decrease and the correlation between the simulated proportions and CRUMBLER estimates also decreased. Nevertheless, by generation 10 44% of animals had their genome proportions estimated with a correlation of at least 70%. In the U.S. commercial crossbreeding does not usually involve the use of more than 3-4 breeds of cattle and while the number of generations of crossbreeding may very well be 10 or perhaps more, many generations will involve the mating of animals with similar genome ancestries and the proportions for each breed will be much greater than present in the generation 10 animals in Fig 11. Consequently, the achieved accuracies are likely to be closer to the generation 3 or 5 results where 99% and 68% of animals, respectively, had their genome proportions estimated with a correlation of greater than 80%.

**Fig. 11.**
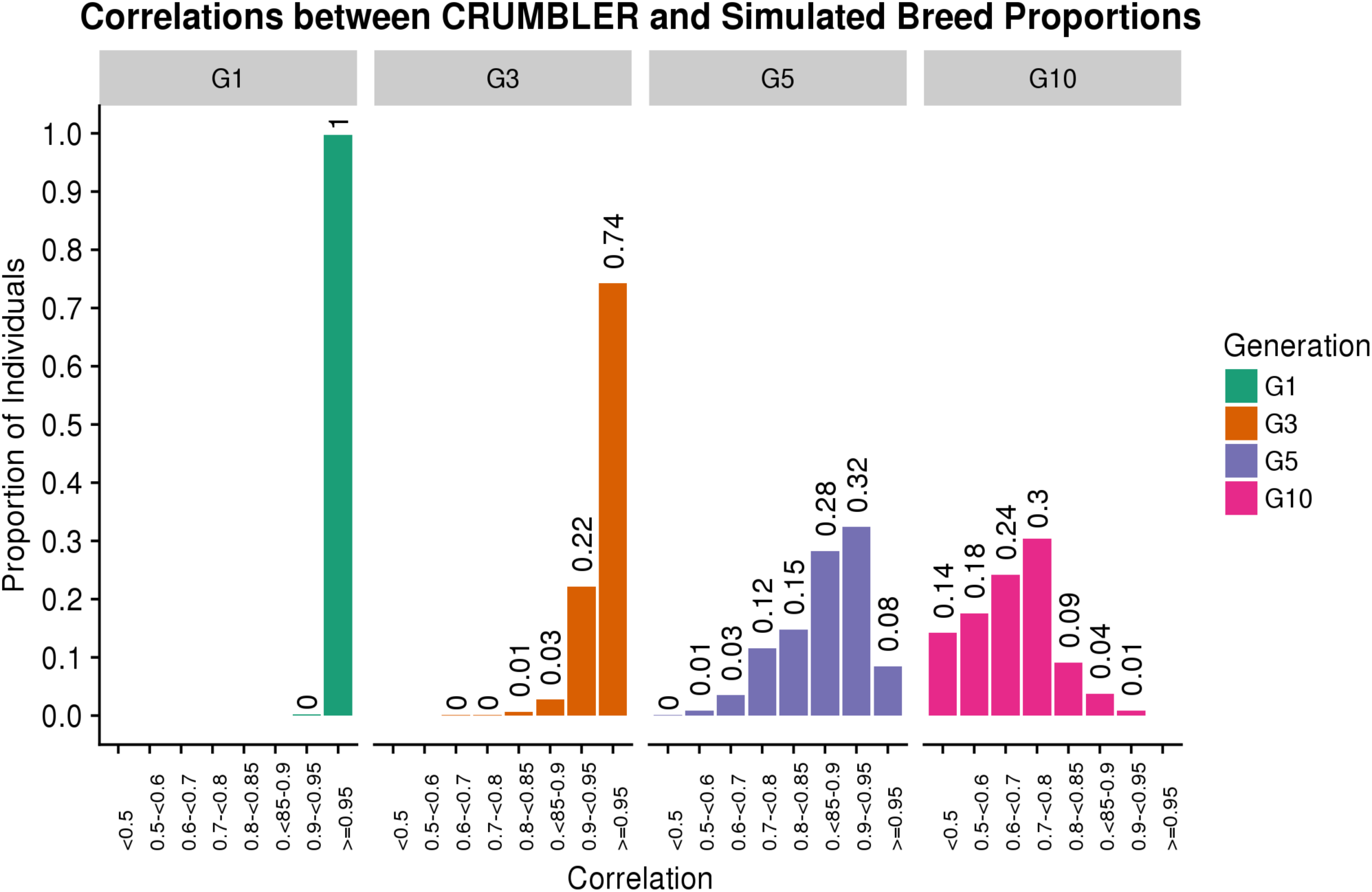
Genotypes were simulated for the indicated number of generations of random mating, with generation 1 (G1) animals being 50:50 proportion except when two parents from the same breed were mated. SNPweights results were obtained using CRUMBLER pipeline parameters correlations between these estimates and the known simulated breed compositions were produced and the proportion of individuals within each correlation class is indicated.

#### Advanced generation composite animals

The ancestry model assumes that neither drift or selection has acted to alter the allele frequencies from those created by the initial admixture proportions. We examined CRUMBLER estimates of breed composition for advanced generation members of the Brangus (n=11,362), Beefmaster (n=3,832) and Santa Gertrudis (n=2,010) composite breeds where selection has had the opportunity to change breed composition from expectations at breed formation. Brangus individuals are expected to be ⅝ Angus and ⅜ Brahman, Beefmaster individuals ¼ Hereford, ¼ Shorthorn and ½ Brahman, and Santa Gertrudis ⅝ Shorthorn and ⅜ Brahman, respectively. These breeds use mating strategies that produce individuals that are expected to possess these proportions for registration within each of the respective breed’s herdbook. However, registerable animals are ultimately advanced generation composites and so drift, meiotic sampling of parental chromosomes and selection are all expected to create individual variation in these ancestry proportions. CRUMBLER results for these advanced generation composites, also known as the American breeds, are shown in Figure 12. Table 6 contains the average breed proportion estimates assigned to each of these breeds by CRUMBLER and their standard deviations across the animals analyzed for each breed. In every instance, CRUMBLER underestimates the expected proportions for each of the American breed populations, however, the ancestral breeds clearly dominate the assignments (Table 6). Interestingly, on average, CRUMBLER estimated proportions of Holstein ancestry for advanced generation Beefmaster and Brangus animals (Figure 12 and Table 6). These American breeds do not contain any Holstein introgression and they do not contain ancestry from a “Ghost Population”, a population that is not present in the reference set, which would lead to a breed assignment to a reference breed that it most closely resembled [6]. We speculate that this effect is caused by selection creating a deviation in allele frequencies from those found in the founder breeds which the model explains by an introgression from a distantly related breed, in this case, Holstein. Stratifying these genotyped animals according to the number of generations from foundation fullblood animals and examining the extent of estimated Holstein introgression, which would be expected to increase with generation number, would enable this to be tested, but we did not have access to the necessary data. However, this hypothesis is supported by the fact that the Santa Gertrudis had the least estimated Holstein introgression and the breed has published estimates of additive genetic merit for many fewer years than the Beefmaster or Brangus.

**Fig. 12.**
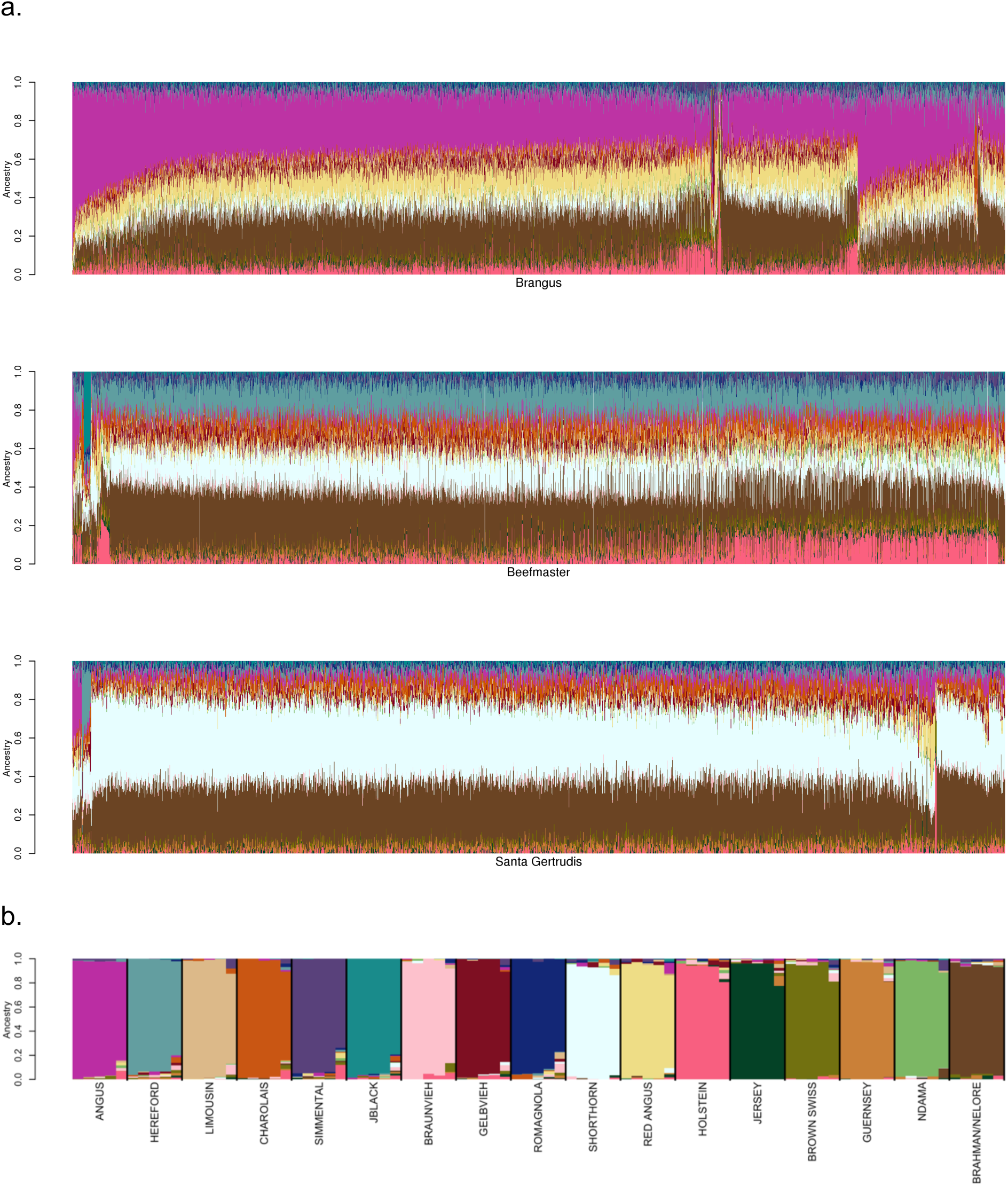
(a) SNPweights ancestry results using CRUMBLER pipeline for 11,362 Brangus, 3,832 Beefmaster, and 2,010 Santa Gertrudis individuals. (b) Breed assignment for these advanced generation composite animals can be determined using this reference breed key.

**Table 6.**
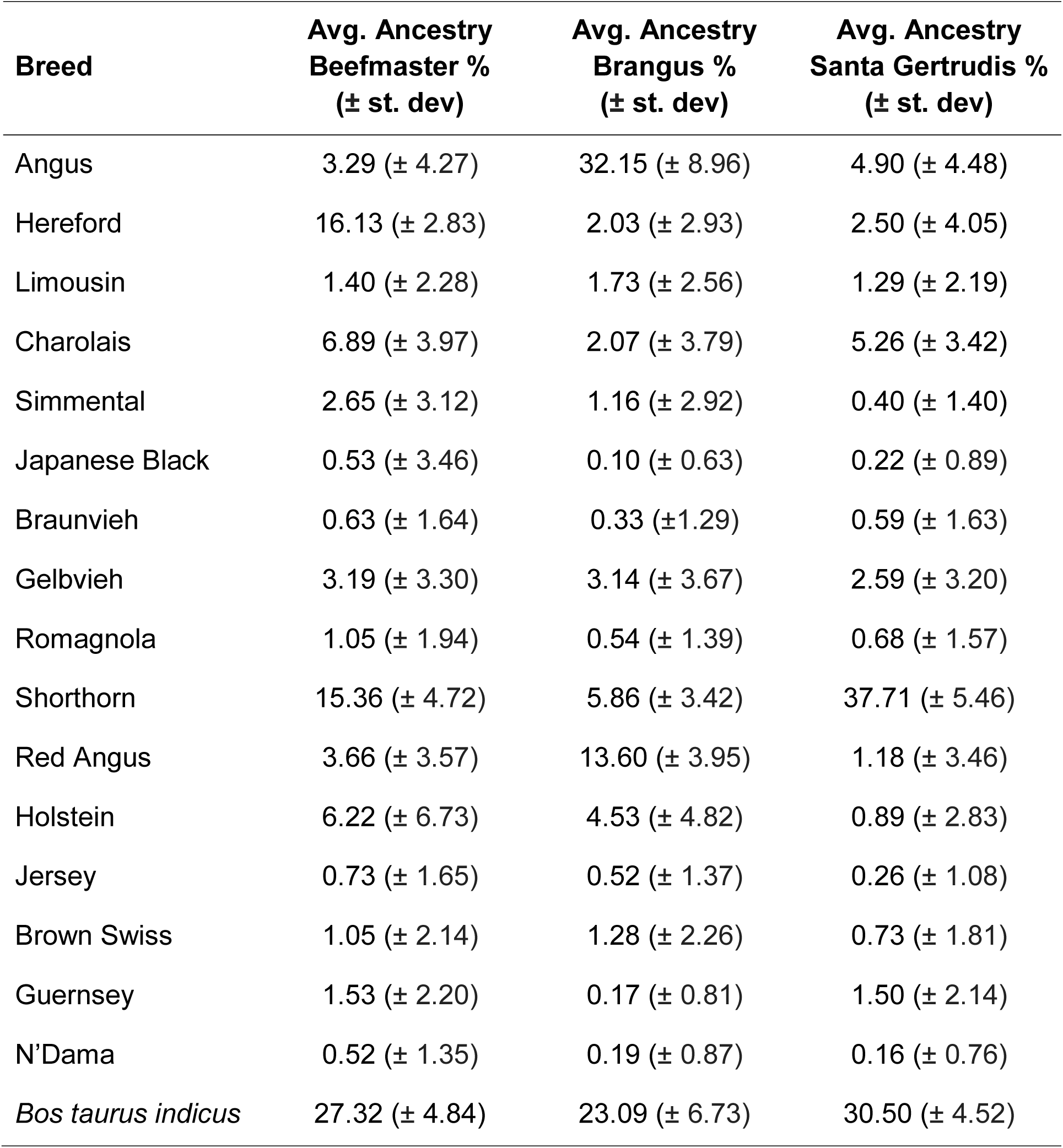
Average breed ancestry percentages assigned to American Breed individuals.

| **Breed** | **Avg. Ancestry**<br>**Beefmaster %**<br>**(± st. dev)** | **Avg. Ancestry**<br>**Brangus %**<br>**(± st. dev)** | **Avg. Ancestry**<br>**Santa Gertrudis %**<br>**(± st. dev)** |
| --- | --- | --- | --- |
| Angus | 3.29 (± 4.27) | 32.15 (± 8.96) | 4.90 (± 4.48) |
| Hereford | 16.13 (± 2.83) | 2.03 (± 2.93) | 2.50 (± 4.05) |
| Limousin | 1.40 (± 2.28) | 1.73 (± 2.56) | 1.29 (± 2.19) |
| Charolais | 6.89 (± 3.97) | 2.07 (± 3.79) | 5.26 (± 3.42) |
| Simmental | 2.65 (± 3.12) | 1.16 (± 2.92) | 0.40 (± 1.40) |
| Japanese Black | 0.53 (± 3.46) | 0.10 (± 0.63) | 0.22 (± 0.89) |
| Braunvieh | 0.63 (± 1.64) | 0.33 (± 1.29) | 0.59 (± 1.63) |
| Gelbvieh | 3.19 (± 3.30) | 3.14 (± 3.67) | 2.59 (± 3.20) |
| Romagnola | 1.05 (± 1.94) | 0.54 (± 1.39) | 0.68 (± 1.57) |
| Shorthorn | 15.36 (± 4.72) | 5.86 (± 3.42) | 37.71 (± 5.46) |
| Red Angus | 3.66 (± 3.57) | 13.60 (± 3.95) | 1.18 (± 3.46) |
| Holstein | 6.22 (± 6.73) | 4.53 (± 4.82) | 0.89 (± 2.83) |
| Jersey | 0.73 (± 1.65) | 0.52 (± 1.37) | 0.26 (± 1.08) |
| Brown Swiss | 1.05 (± 2.14) | 1.28 (± 2.26) | 0.73 (± 1.81) |
| Guernsey | 1.53 (± 2.20) | 0.17 (± 0.81) | 1.50 (± 2.14) |
| N'Dama | 0.52 (± 1.35) | 0.19 (± 0.87) | 0.16 (± 0.76) |
| *Bos taurus indicus* | 27.32 (± 4.84) | 23.09 (± 6.73) | 30.50 (± 4.52) |

### Admixture

We also tested the ADMIXTURE software [22] for ancestry estimation and integration into the CRUMBLER pipeline using the same reference breed panel that was developed for use with SNPweights. ADMIXTURE uses maximum likelihood estimation to fit the same statistical model as STRUCTURE, however, STRUCTURE does not allow the specification of individuals of known descent to be used as a reference panel [22]. ADMIXTURE allows a supervised analysis, in which the user can specify a reference set of individuals, by specifying the “--supervised” flag and requires an additional file with a “.pop” suffix to specify the genotypes of the reference population individuals [22]. Unlike SNPweights, the reference population individuals’ genotypes must be provided in a genotype file for each analysis.

We first conducted an ADMIXTURE analysis in which we self-assigned ancestry for the animals in the reference breed set formed with ≤50 individuals per breed from the individuals that had ≥85% assignment to the breed of registration (Fig. 13). The results shown in Fig. 13 are similar to those in Fig. 6 for the same reference panel, albeit with perhaps less evidence of background introgression. We next conducted an analysis using the reference panel used in Fig. 13 merged with data for the 2,005 high percentage crossbred Herefords animals. The results shown in Fig. 14, reveal a significant change in the ancestry proportions estimated for the reference panel Guernsey, Gelbvieh and Romagnola individuals between the two analyses which used exactly the same reference panel, but differed only in the number of individuals for which ancestry was to be estimated. This suggests that ADMIXTURE may use the target individuals to update information provided by the reference panel individuals specified in the “.pop” file. Consequently, the ADMIXTURE estimated ancestry proportions appear to be context dependent and may vary based on the other individuals included in the analysis.

**Fig. 13.**
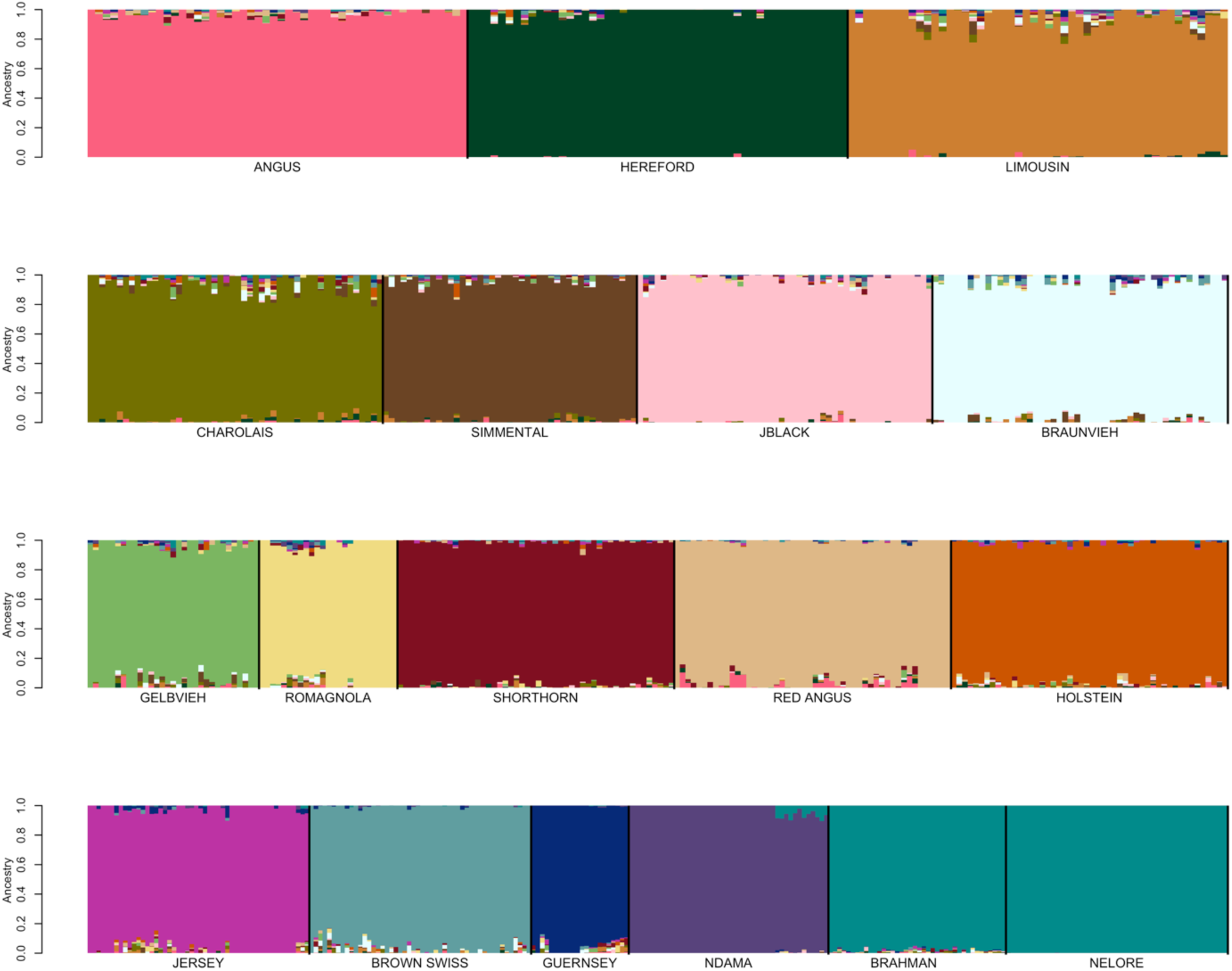
Self-assignment of ancestry for the animals in the reference breed set formed with ≤50 individuals per breed from the individuals that had ≥85% assignment to their breed of registration using ADMIXTURE.

**Fig. 14.**
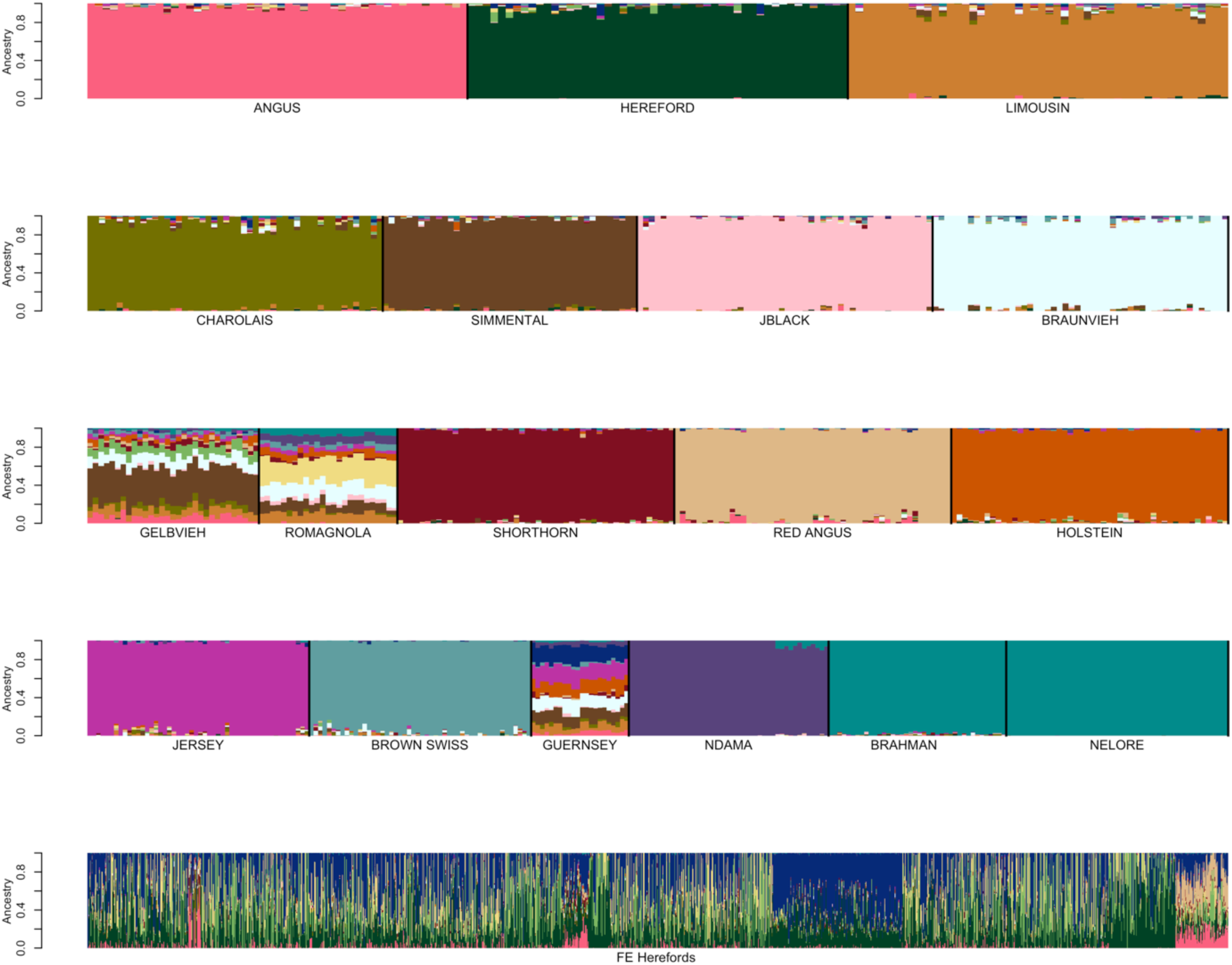
ADMIXTURE analysis conducted using the same data as shown in Figure 13 (first four rows), merged with an additional 2,005 high percentage crossbred Hereford target individuals (last row). Here, the 2,005 Hereford crossbred individuals appear after the reference individuals in the input genotype file.

Moreover, the order in which the target individuals appear in the genotype input file also appears to affect ADMIXTURE estimates of ancestry proportions for the target individuals. Fig. 15 shows the results of an ADMIXTURE analysis in which the target individuals were identical to those shown in Fig. 14, but for which the order of the reference individuals and the 2,005 Hereford crossbred individuals was reversed in the input files. In Fig. 14, the reference individuals appear before the 2,005 Hereford crossbred individuals in the input file, whereas in Fig. 15, the 2,005 Hereford crossbred individuals appeared before the reference individuals in the input file. The results reveal a significant change in ancestry proportions for Guernsey and Gelbvieh, but the Romagnola now appear to be non-admixed. Finally, we performed an ADMIXTURE analysis for these animals in which the order of animals in the input genotype file was completely randomized (Fig. 16). Following analysis, the individuals were sorted to generate Fig. 16. Again, the ancestry proportions for the Guernsey, Gelbvieh and Romagnola individuals suggest these breeds to be admixed.

**Fig. 15.**
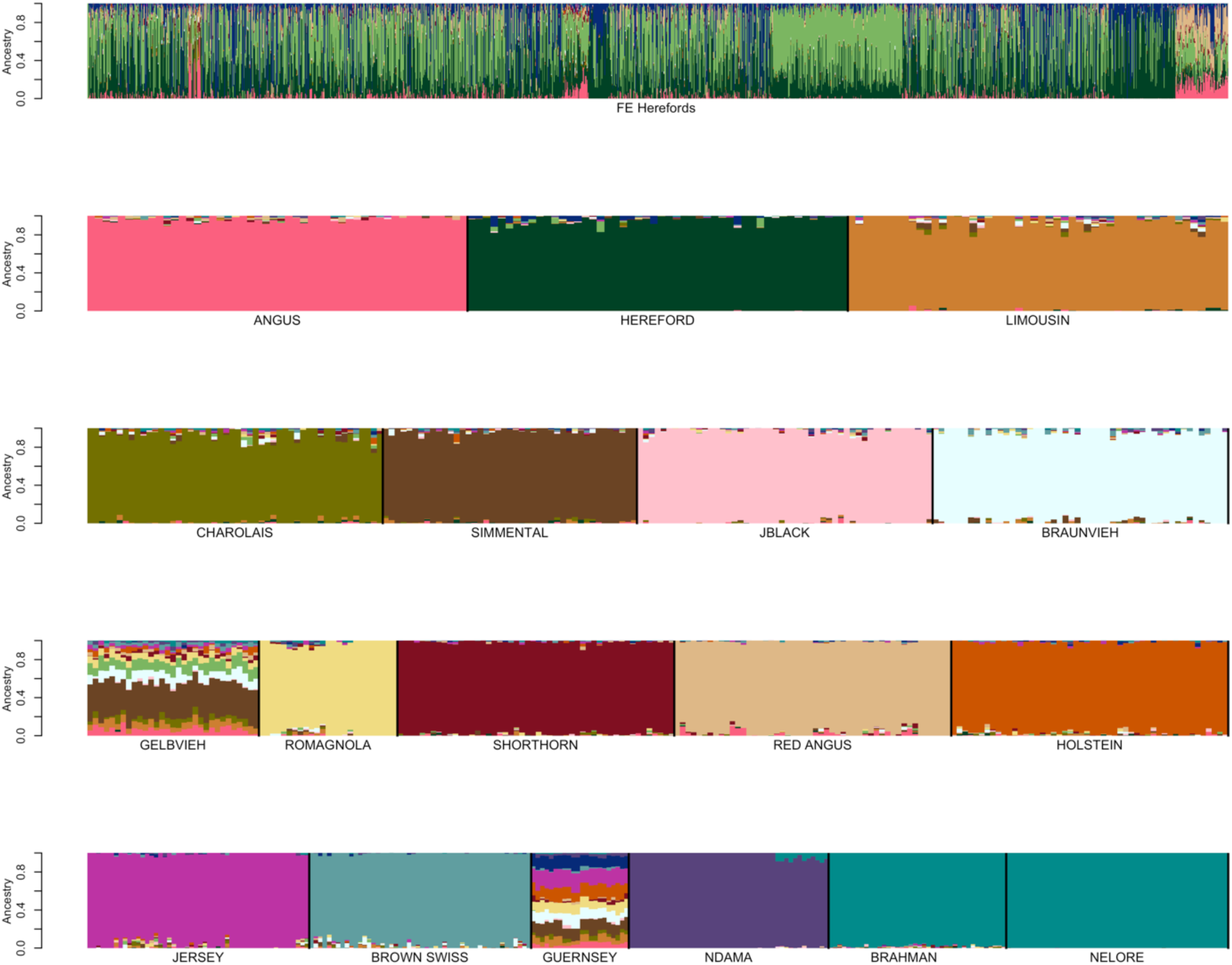
ADMIXTURE analysis conducted using the same data as shown in Fig. 14. Here, the 2,005 Hereford crossbred individuals appear before the reference individuals in the input genotype file. The first row represents the 2005 Hereford crossbred samples. Rows 2 to 5 show the reference panel individuals.

**Fig. 16.**
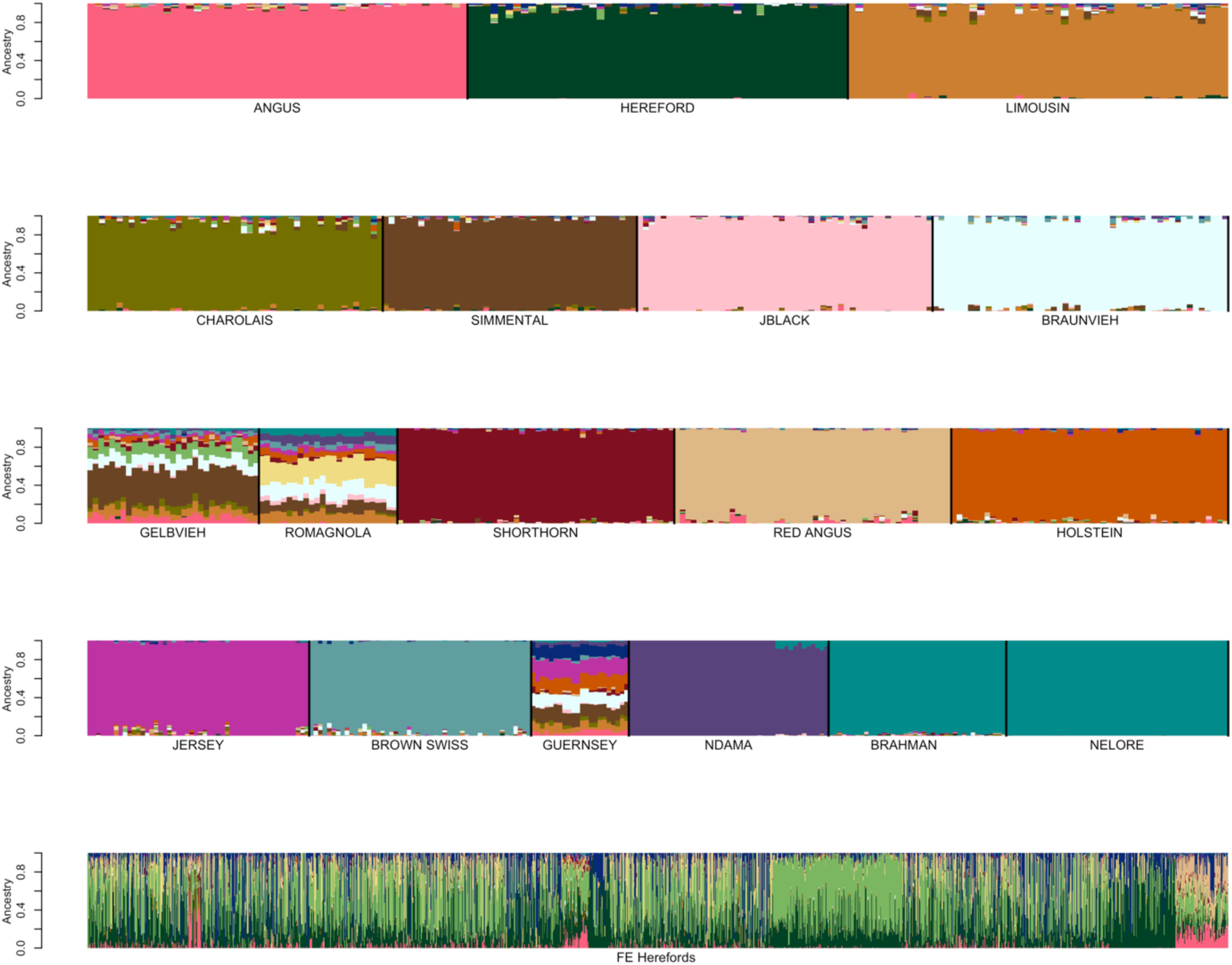
ADMIXTURE analysis conducted using the same data as shown in Figs. 14 and 15, but with the order of the individuals in the input genotype file randomized. The animals were sorted following analyses to generate this figure where the first four rows represent the reference panel individuals, the fifth row shows the 2,005 Hereford crossbred animals.

STRUCTURE and ADMIXTURE are widely used for characterizing admixed populations [6], however, we have not found any reports in the literature that indicate that the software is sensitive to the input order of individuals. However, we suspect that the majority of users would have no need or motivation to run the software with permuted data input files. Nevertheless, because of these inconsistencies between results, we chose to not use ADMIXTURE for ancestry estimation within the CRUMBLER pipeline.

### Broader application using additional commercially available assays

To broaden the spectrum of data from different commercially available assays that can be evaluated, an additional intersection of markers was obtained using 11 commercially available bovine assays including the GGP-90KT, GGP-F250, GGP-HDV3, GGP-LDV3, GGP-LDV4, BovineHD, BovineSNP50, i50K, Irish Cattle Breeding Federation (Cork, Ireland) IDBv3, and GeneSeek (Lincoln, NE) BOVG50v1 assays. The intersection SNP set included 6,363 SNPs (BC6K). A SNPweights self-assignment analysis using the reference set of individuals with ≥85% assignment to their breed of registration was conducted to assess the effects of the reduction in number of markers used for ancestry assignment. The ancestry proportions assigned based on the BC6K marker set (Fig. 17) did not differ appreciably from those obtained using the BC7K marker set (Fig. 6). This result indicates the utility of CRUMBLER and the reference panel breed set across the spectrum of commercially available genotyping platforms.

**Fig. 17.**
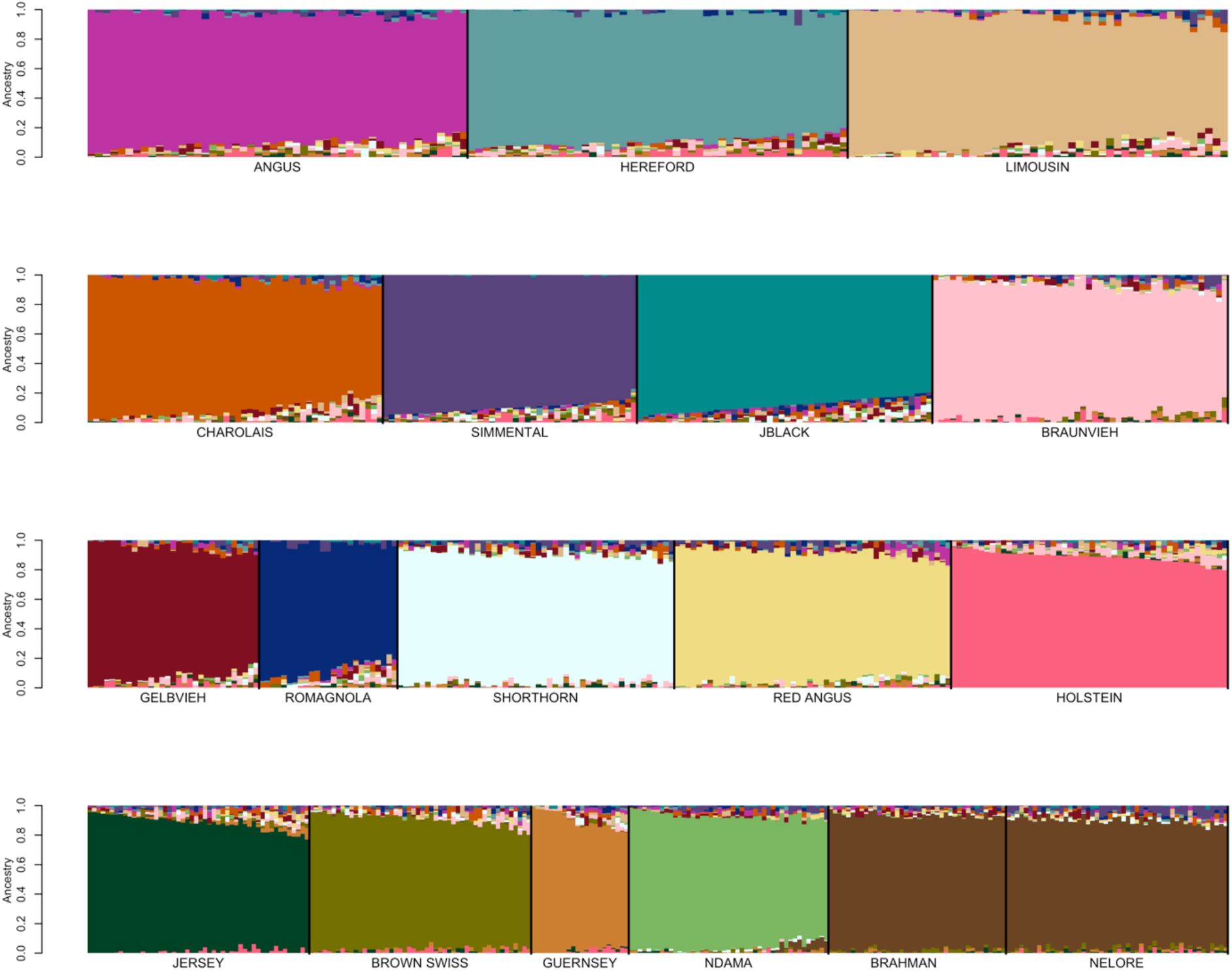
Reference breed panel constructed by the random sampling of ≤50 individuals per breed from individuals with ≥85% ancestry was self-assigned to reference breed ancestry using the BC6K marker set.

## Conclusions

The determination of a set of reference population breeds and individuals that define allele and genotype frequencies at each variant for each of the breeds is arguably the most important, yet technically difficult step in the process of ancestry estimation. We employed several iterations of filtering to remove recently admixed individuals and identify a relatively homogeneous set of individuals that nevertheless represented the variation that might be expected among individuals within a breed. Once determined, the reference panel genotype data need only be processed once to obtain SNP weights removing the need to share genotype data for reference individuals in subsequent studies [11]. The upfront development of an external reference breed panel capitalizes on the rich ancestry information available in large available datasets, and relatedness, variation in sample sizes and diversity among the target individuals does not affect the inference of ancestry [11].

In cattle, the visual evaluation of breed characteristics is a poor method for evaluating the ancestry of individuals. Breed association pedigrees can be used to estimate expected breed compositions, however, the random assortment of chromosomes into gametes and selection can lead to ancestry proportions that differ from those expected based upon pedigree. Moreover, the vast majority of commercial beef cattle in the U.S. have no or very limited pedigree information and since these animals are frequently used for genomic research [3–5], there is a need for a tool that can routinely provide ancestry estimates for downstream use in GWAA or other genetic studies.

We tested ADMIXTURE and SNPweights and found that results from ADMIXTURE appear to depend on the ancestry and order of appearance of individuals within the genotype input file. We therefore developed an analysis pipeline, CRUMBLER, based upon PLINK, EIGENSOFT and SNPweights to automate the process of ancestry estimation. The developed bovine pipeline utilizes the 6,799 SNPs present on 8 commercially utilized bovine SNP genotyping assays and results using these SNPs are consistent with results obtained when 13,291 SNPs were used. From an available 48,776 genotyped individuals, we also developed a reference panel of 806 individuals sampled from 17 breeds to have ≤50 individuals per breed that had ≥85% assignment to their breed of registration. This panel appears to allow the robust estimation of the ancestry of advanced generation admixed animals, however, all breeds share some common ancestry which predates the recent development of breed association herdbooks [16,23].

CRUMBLER is not limited to application in cattle and with the provision of suitable reference breed allele frequencies can be applied to other species for ancestry estimation. CRUMBLER pipeline scripts and reference panel breed SNP weights are available on GitHub (https://github.com/tamarcrum/CRUMBLER).

## Additional files

**Supplementary Information (PDF).** This file contains the source code changes in SMARTPCA within versions of EIGENSOFT beyond 5.0.2 to enable compatibility with SNPweights.

**Supplementary Methods (PDF).** This file describes the preliminary fastSTRUCTURE analyses conducted on subsamples of breeds in the development of the reference breed panel.

**Supplementary Figures (PDF).** This file contains Supplementary Figures S1-S16.

Fig. S1 An overview of the processes and iterations of filtering conducted in the development of the reference panel.

Fig. S2 Preliminary FastSTRUCTURE analysis of candidate Angus and Simmental reference population animals.

Fig. S3 Preliminary fastSTRUCTURE analysis of candidate Angus and Gelbvieh reference population animals.

Fig. S4 Preliminary fastSTRUCTURE analysis of candidate Angus and Limousin reference population animals.

Fig. S5 Preliminary fastSTRUCTURE analysis of candidate Angus and Red Angus reference population animals.

Fig. S6 Preliminary fastSTRUCTURE analysis of candidate Red Angus, Hereford, Shorthorn and Salers reference population animals.

Fig. S7 Preliminary fastSTRUCTURE analysis of candidate Red Angus, Hereford and Shorthorn reference population animals.

Fig. S8 Preliminary fastSTRUCTURE analysis of candidate N’Dama, Nelore and Brahman reference population animals.

Fig. S9 SNPweights self-assignment analysis for the reference sample set containing ≤200 individuals per breed analyzed using the BC7K marker set.

Fig. S10 SNPweights self-assignment analysis for the reference sample set containing ≤150 individuals per breed analyzed using the BC7K marker set.

Fig. S11 SNPweights self-assignment analysis for the reference sample set containing ≤50 individuals per breed analyzed using the BC7K marker set.

Fig. S12 SNPweights self-assignment analysis for the reference sample sets containing ≤50 individuals per breed analyzed using the BC13K marker set.

Fig. S13 SNPweights self-assignment analysis for the reference sample set with ≥80% ancestry to breed of registry and ≤50 individuals per breed using the BC7K marker set.

Fig. S14 SNPweights self-assignment analysis for reference sample set with ≥75% ancestry to breed of registry and ≤50 individuals per breed using the BC7K marker set.

Fig. S15 SNPweights self-assignment analysis for the reference sample set with ≥70% ancestry to breed of registry and ≤50 individuals per breed using the BC7K marker set.

Fig. S16 SNPweights self-assignment analyses using a reference panel with ≤50 individuals per breed and sampling from the individuals with ≥85% assignment to their breed of registry but with (a) Red Angus or (b) Angus excluded from the reference panel.

## Authors’ contributions

TC conceived the study and managed the project. TC, RS, JD, and JT contributed to defining the research questions and analytical approaches and interpretation of the results. TC programmed the CRUMBLER pipeline and carried out the data analyses. TC and JT drafted the manuscript. LR provided the Nelore samples but did not have involvement in the scientific direction. All authors read and approved the final manuscript.

## Availability of data and materials

Project Name: CRUMBLER

Project Home Page: https://github.com/tamarcrum/CRUMBLER

Programming Language: Python

Other Requirements: PLINK, EIGENSOFT, and SNPweights

License: GNU GPL

## Ethics approval and consent to participate

Not applicable.

## Consent for Publication

Not applicable.

## Acknowledgements

Not applicable.

