## Supporting Figures for "CRUMBLER: A Tool for the Prediction of Ancestry in Cattle"

**Supplementary Figures.** (PDF)

**Fig. S1 An overview of the processes and iterations of filtering conducted in the development of the reference panel.**

**Fig. S2 Preliminary FastSTRUCTURE analysis of candidate Angus and Simmental reference population animals.**

**Fig. S3 Preliminary fastSTRUCTURE analysis of candidate Angus and Gelbvieh reference population animals.**

**Fig. S4 Preliminary fastSTRUCTURE analysis of candidate Angus and Limousin reference population animals.**

**Fig. S5 Preliminary fastSTRUCTURE analysis of candidate Angus and Red Angus reference population animals.**

**Fig. S6 Preliminary fastSTRUCTURE analysis of candidate Red Angus, Hereford, Shorthorn and Salers reference population animals.**

**Fig. S7 Preliminary fastSTRUCTURE analysis of candidate Red Angus, Hereford and Shorthorn reference population animals.**

**Fig. S8 Preliminary fastSTRUCTURE analysis of candidate N'Dama, Nelore and Brahman reference population animals.**

**Fig. S9 SNPweights self-assignment analysis for the reference sample set containing ≤200 individuals per breed analyzed using the BC7K marker set.**

**Fig. S10 SNPweights self-assignment analysis for the reference sample set containing ≤150 individuals per breed analyzed using the BC7K marker set.**

**Fig. S11 SNPweights self-assignment analysis for the reference sample set containing ≤50 individuals per breed analyzed using the BC7K marker set.**

**Fig. S12 SNPweights self-assignment analysis for the reference sample sets containing ≤50 individuals per breed analyzed using the BC13K marker set.**

**Fig. S13 SNPweights self-assignment analysis for the reference sample set with ≥80% ancestry to breed of registry and ≤50 individuals per breed using the BC7K marker set.**

**Fig. S14 SNPweights self-assignment analysis for reference sample set with ≥75% ancestry to breed of registry and ≤50 individuals per breed using the BC7K marker set.**

**Fig. S15 SNPweights self-assignment analysis for the reference sample set with ≥70% ancestry to breed of registry and ≤50 individuals per breed using the BC7K marker set.**

**Fig. S16 SNPweights self-assignment analyses using a reference panel with ≤50 individuals per breed and sampling from the individuals with ≥85% assignment to their breed of registry but with (a) Red Angus or (b) Angus excluded from the reference panel.**

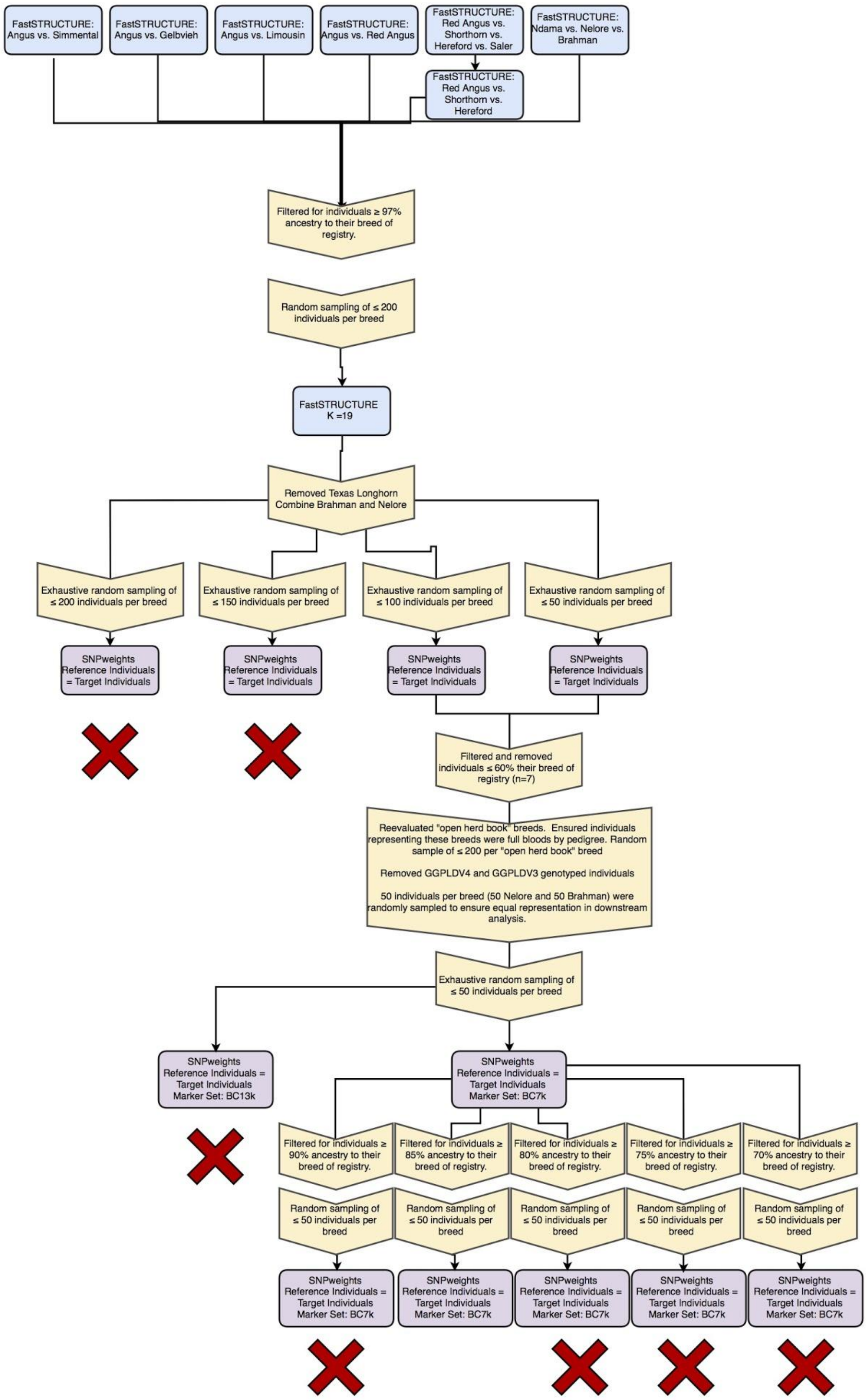


**Fig. S1** An overview of the processes and iterations of filtering conducted in the development of the reference panel. Blue = FastSTRUCTURE analyses, Purple = SNPweights analyses, Yellow Arrows = Data management processes.

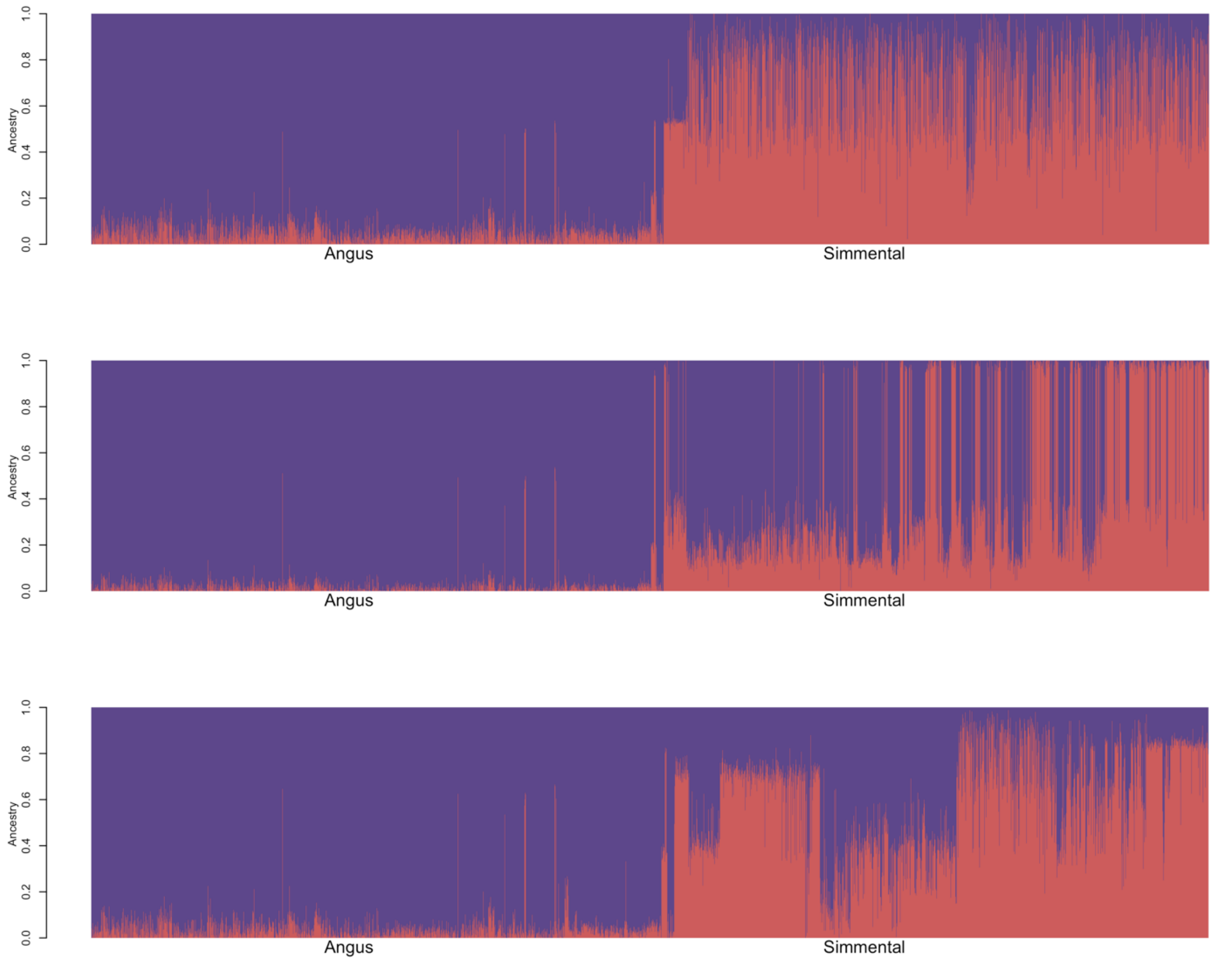


**Fig. S2** Preliminary FastSTRUCTURE analysis of candidate Angus and Simmental reference population animals. Three analyses were run to compare equal numbers of animals from each breed when the available sample sizes for each breed differed. Each row represents an analysis and each animal is represented as a vertical line.

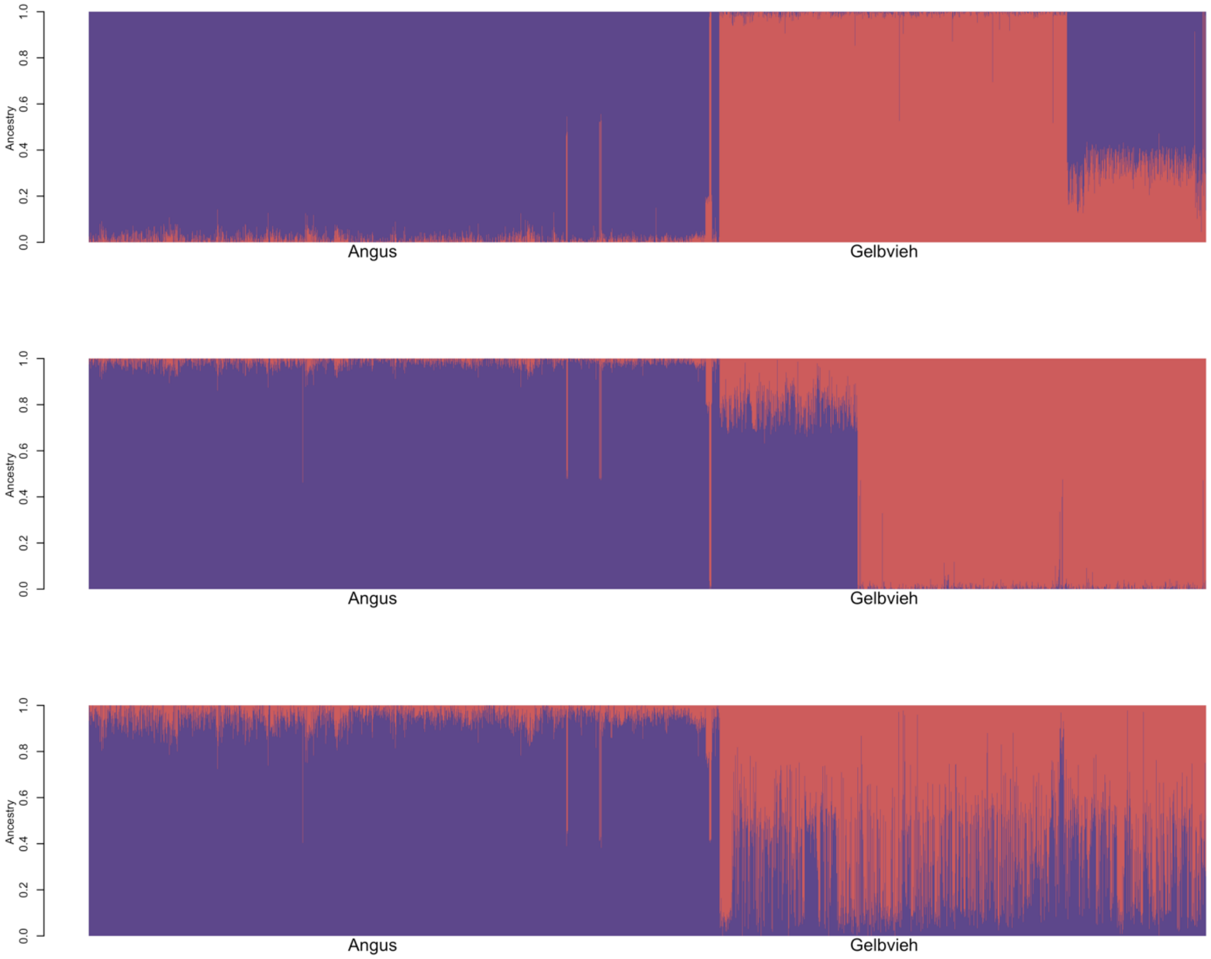


**Fig. S3** Preliminary FastSTRUCTURE analysis of candidate Angus and Gelbvieh reference population animals. Three analyses were run to compare equal numbers of animals from each breed when the available sample sizes for each breed differed. Each row represents an analysis and each animal is represented as a vertical line.

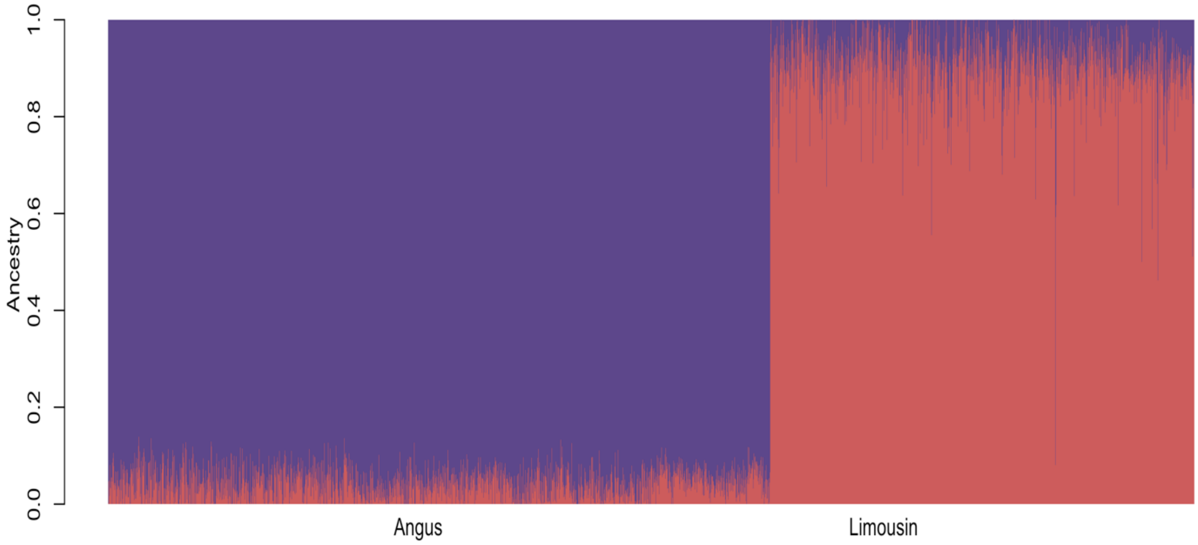


**Fig. S4** Preliminary FastSTRUCTURE analysis of candidate Angus and Limousin reference population animals. Each animal is represented as a vertical line.

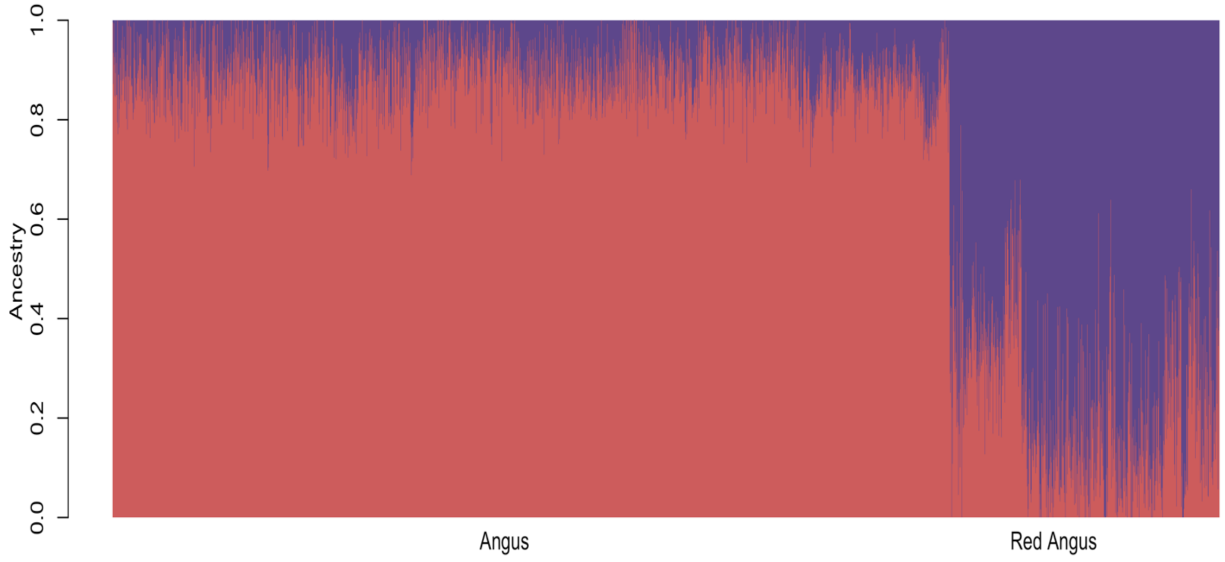


**Fig. S5** Preliminary FastSTRUCTURE analysis of candidate Angus and Red Angus reference population animals. Each animal is represented as a vertical line.

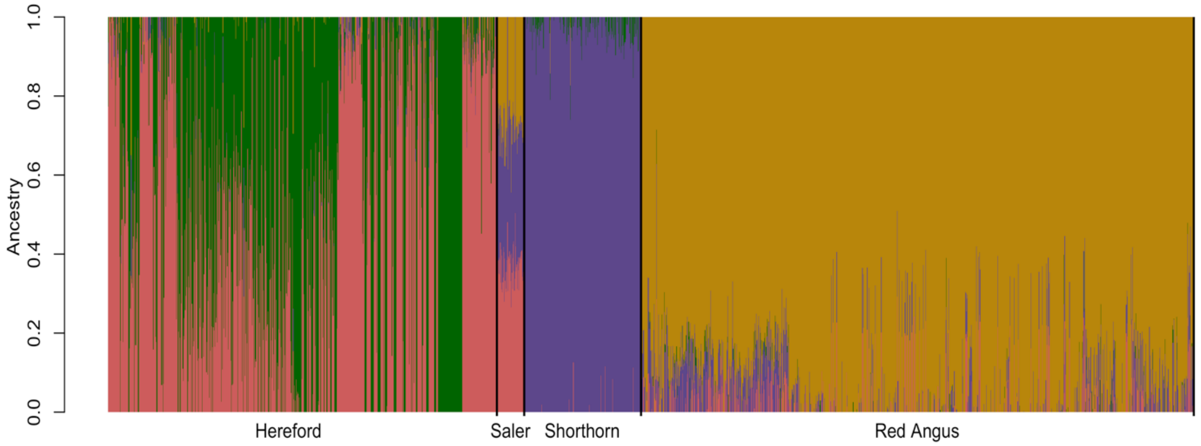


**Fig. S6** Preliminary FastSTRUCTURE analysis of candidate Red Angus, Hereford, Shorthorn, and Salers reference population animals. Each animal is represented as a vertical line.

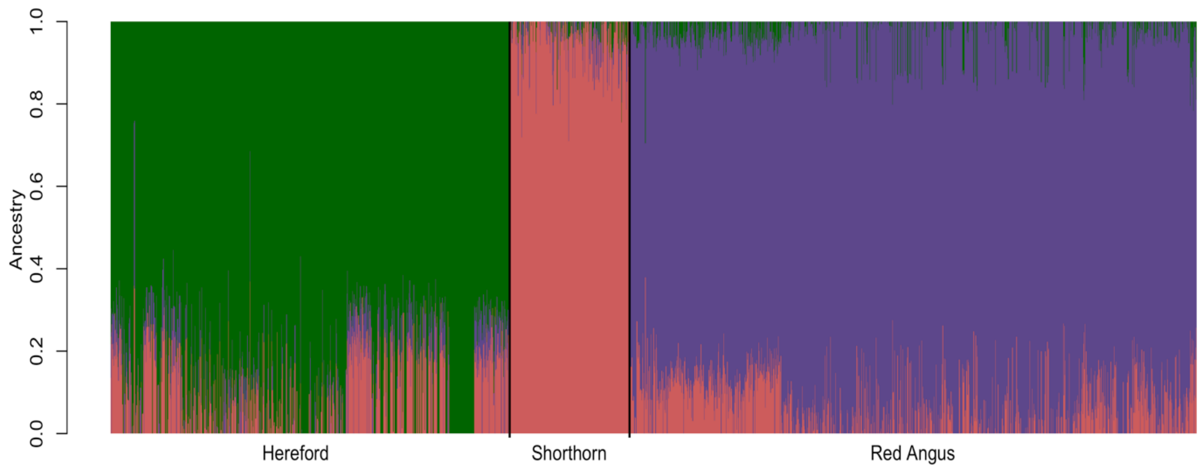


**Fig. S7** Preliminary FastSTRUCTURE analysis of candidate Red Angus, Hereford, and Shorthorn reference population individuals. Each animal is represented as a vertical line.

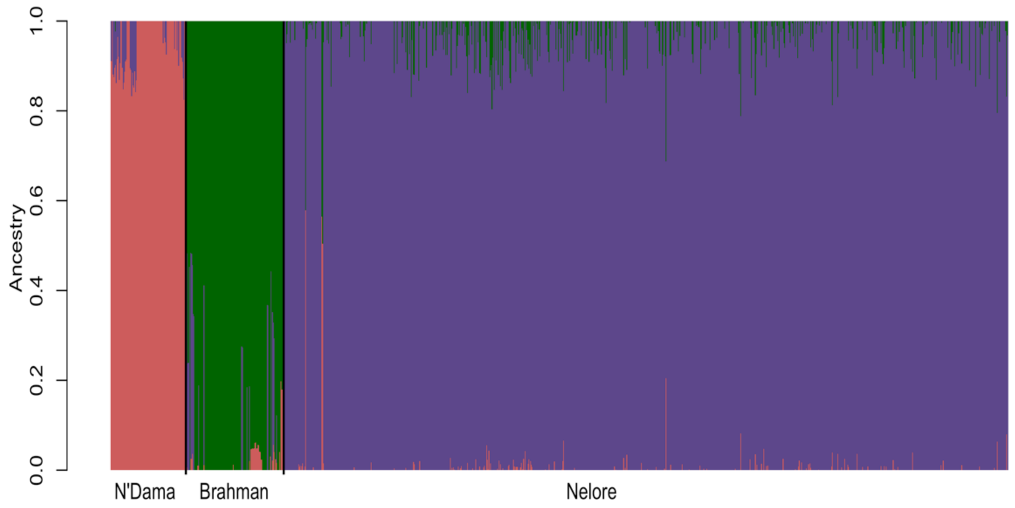


**Fig. S8** FastSTRUCTURE analysis of candidate N'Dama, Brahman and Nelore reference population individuals. Each animal is represented as a vertical line.

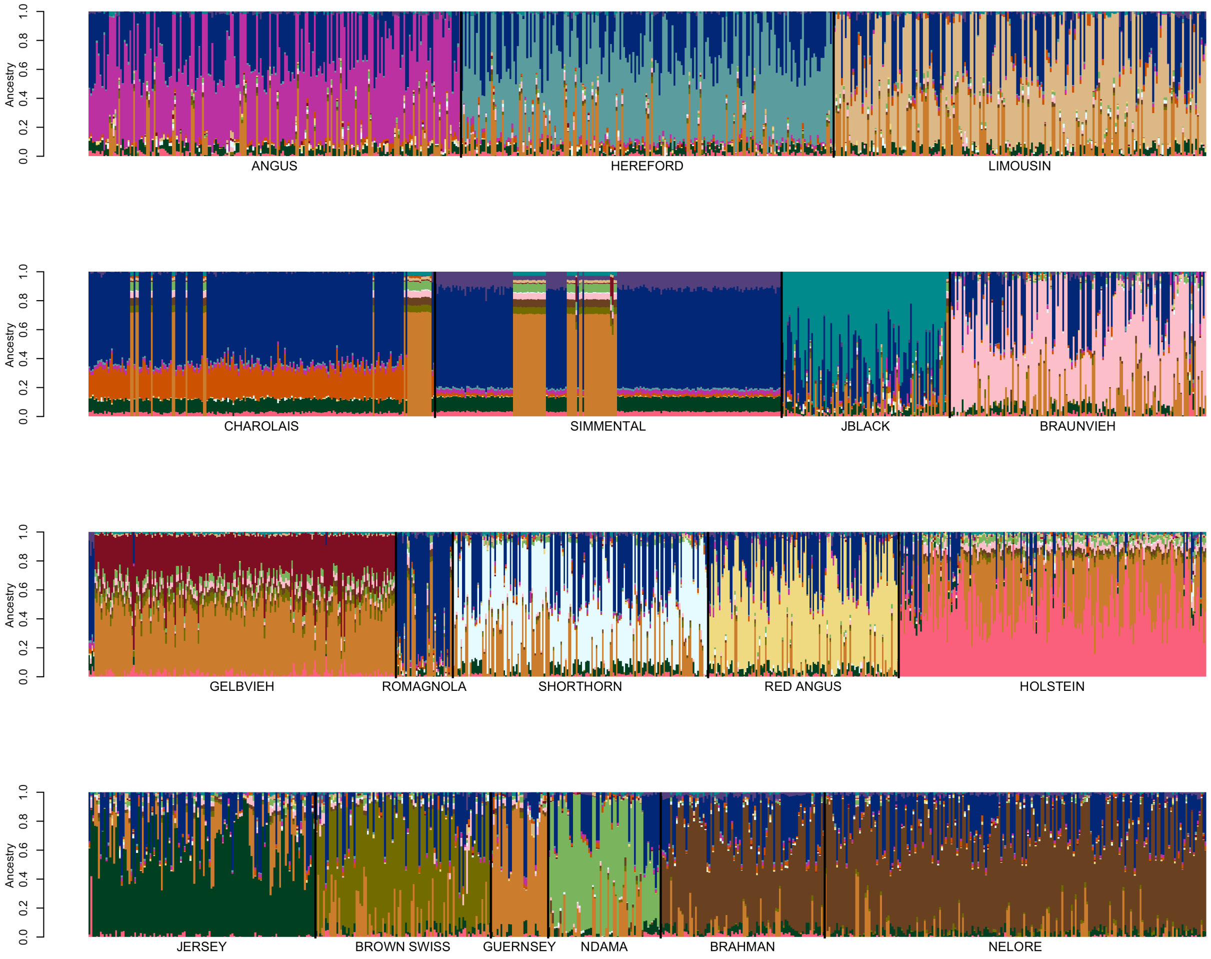


**Fig. S9** SNPweights self-assignment analysis for the reference sample set containing ≤200 individuals per breed analyzed using the BC7K marker set.

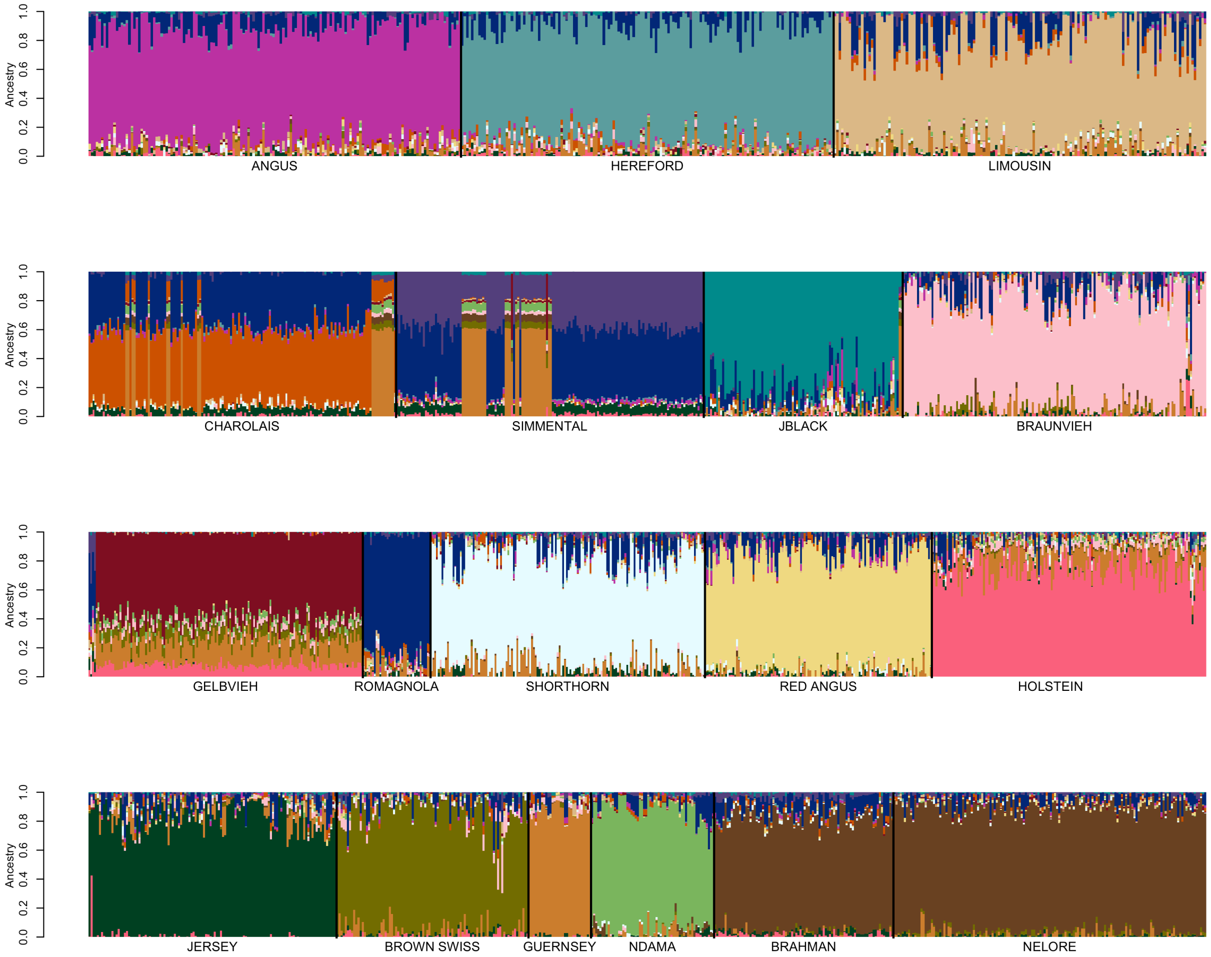


**Fig. S10** SNPweights self-assignment analysis for the reference sample set containing ≤150 individuals per breed analyzed using the BC7K marker set.

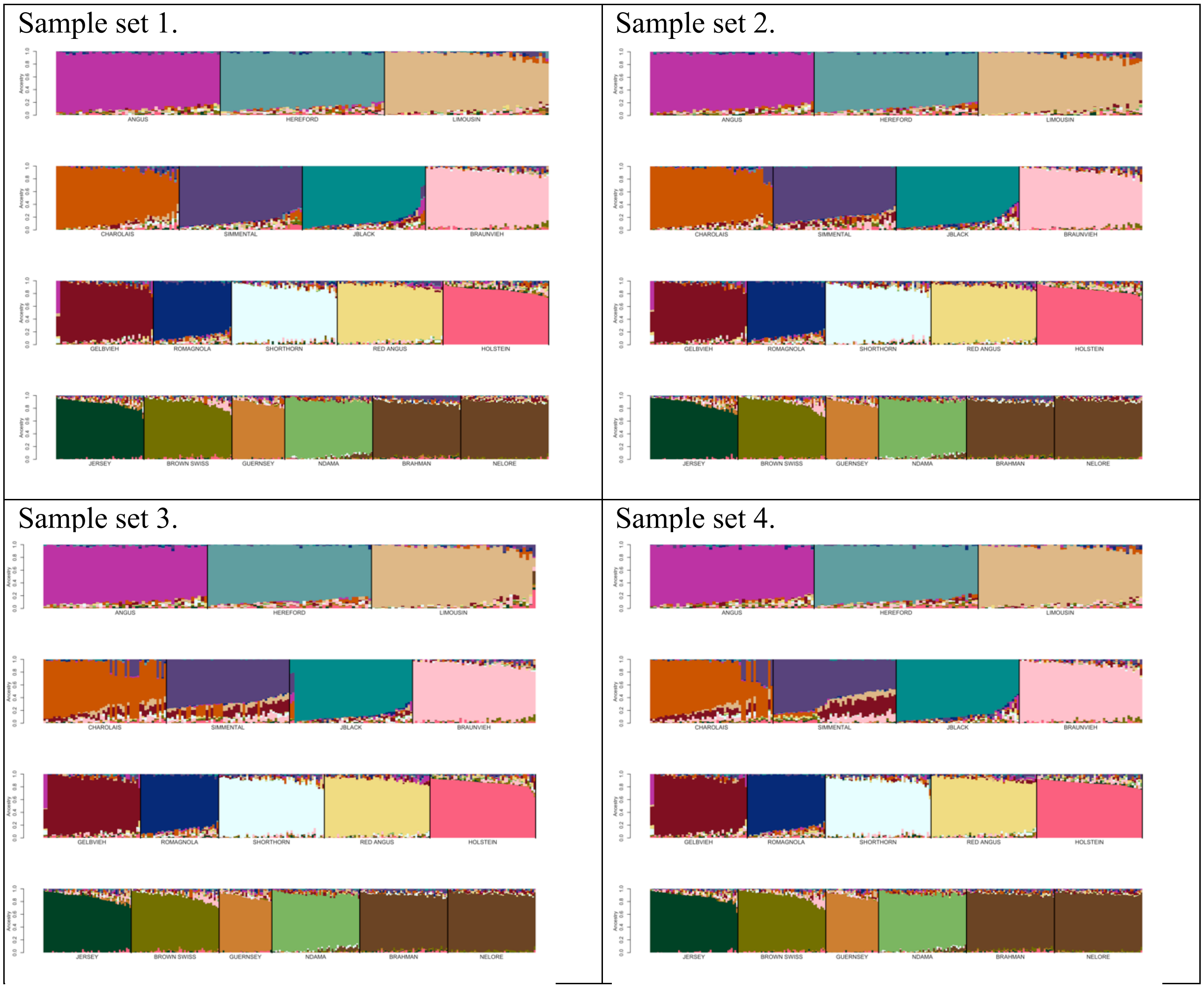


**Fig. S11** SNPweights self-assignment analysis for the reference sample set containing ≤50 individuals per breed analyzed using the BC7K marker set. Reference breed panels were constructed by randomly sampling ≤50 individuals per breed until all individuals were represented in at least one set, resulting in 5 candidate reference sample sets (sample set 5 is shown in Fig. 4a).

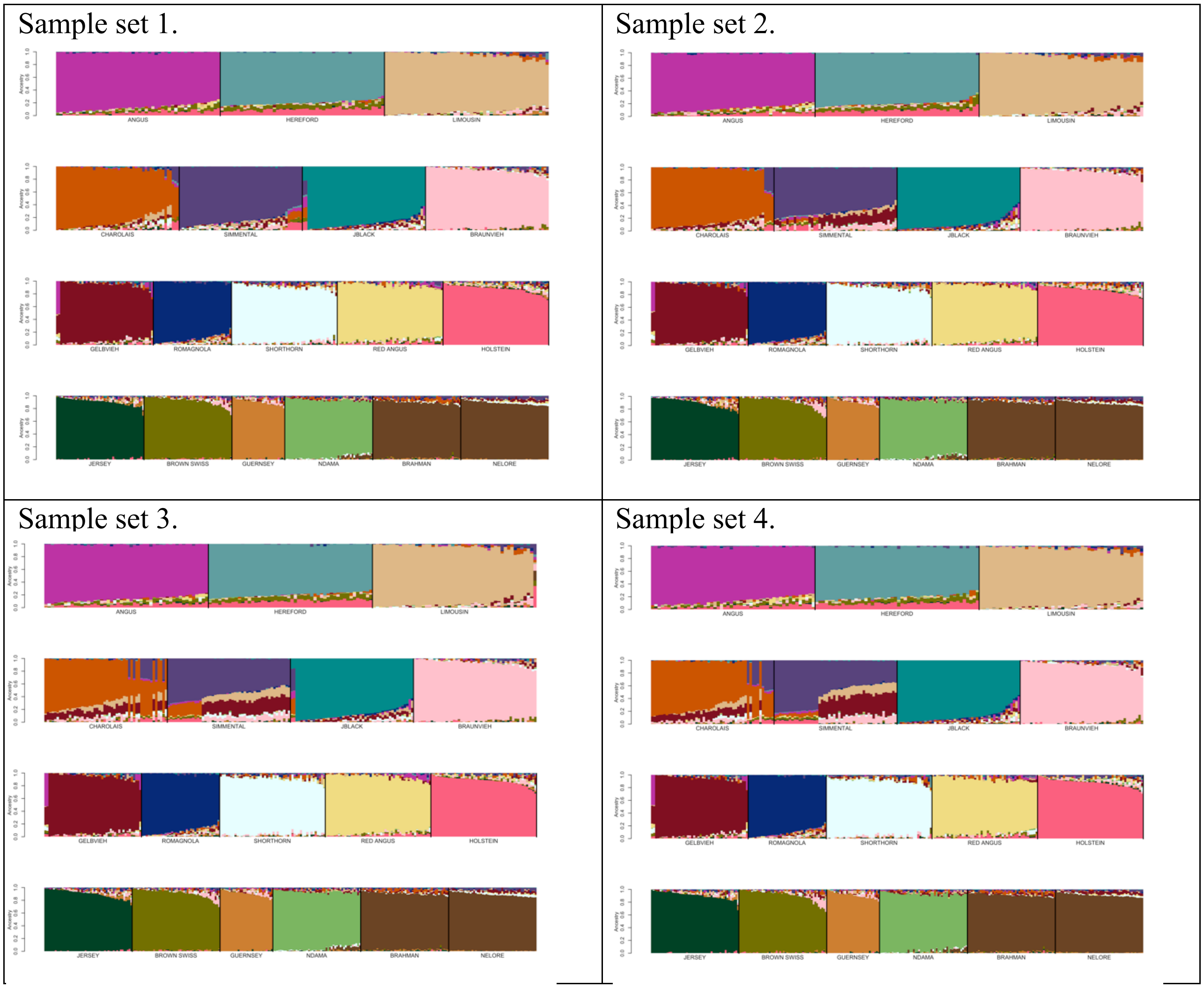


**Fig. S12** SNPweights self-assignment analysis for the reference sample set containing ≤50 individuals per breed analyzed using the BC13K marker set. Reference breed panels were constructed by randomly sampling ≤50 individuals per breed until all individuals were represented in at least one set, resulting in 5 candidate reference sample sets (sample set 5 is shown in Fig. 4b).

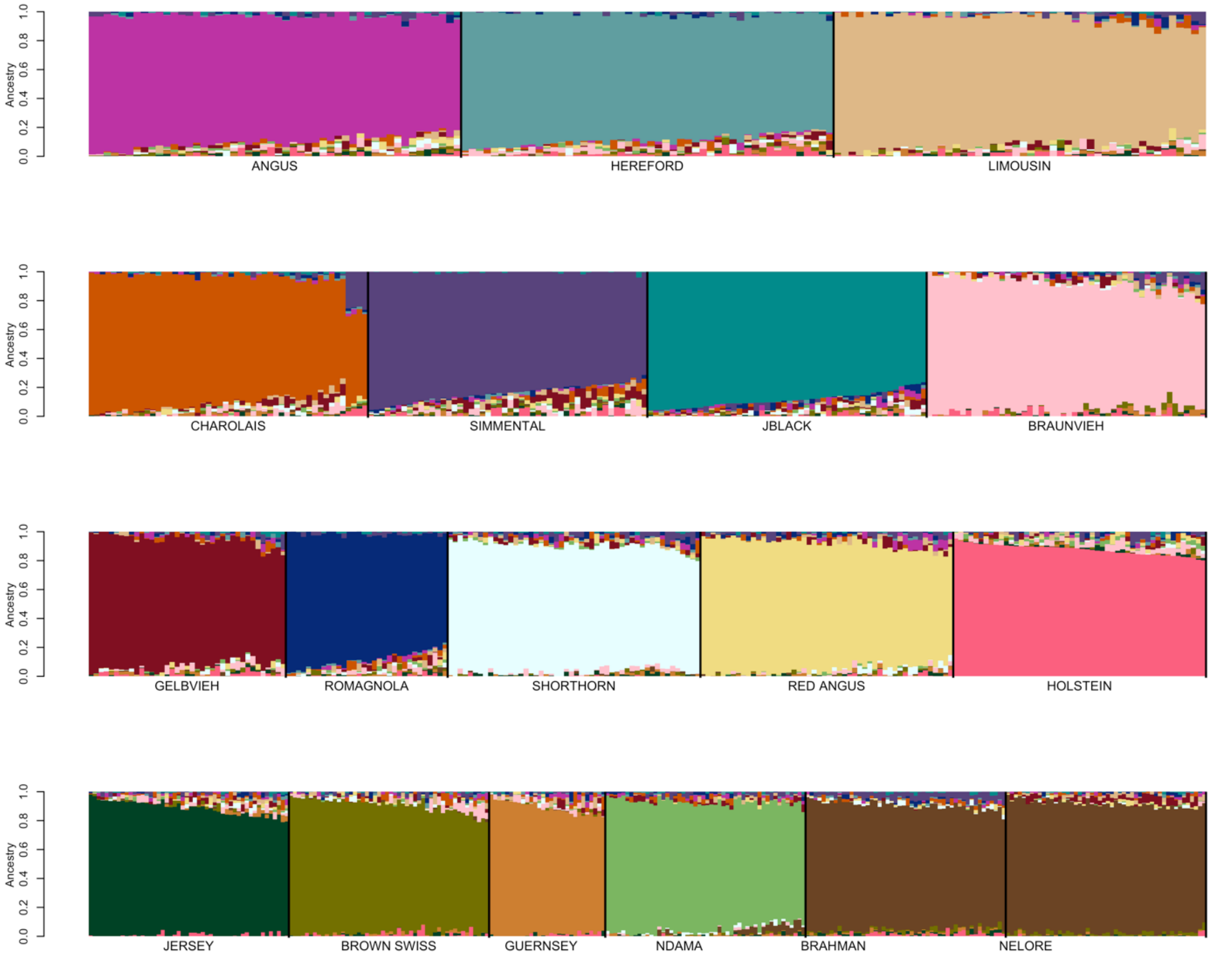


**Fig. S13** SNPweights self-assignment analysis for the reference sample set with ≥80% ancestry to breed of registry and ≤50 individuals per breed using the BC7K marker set.

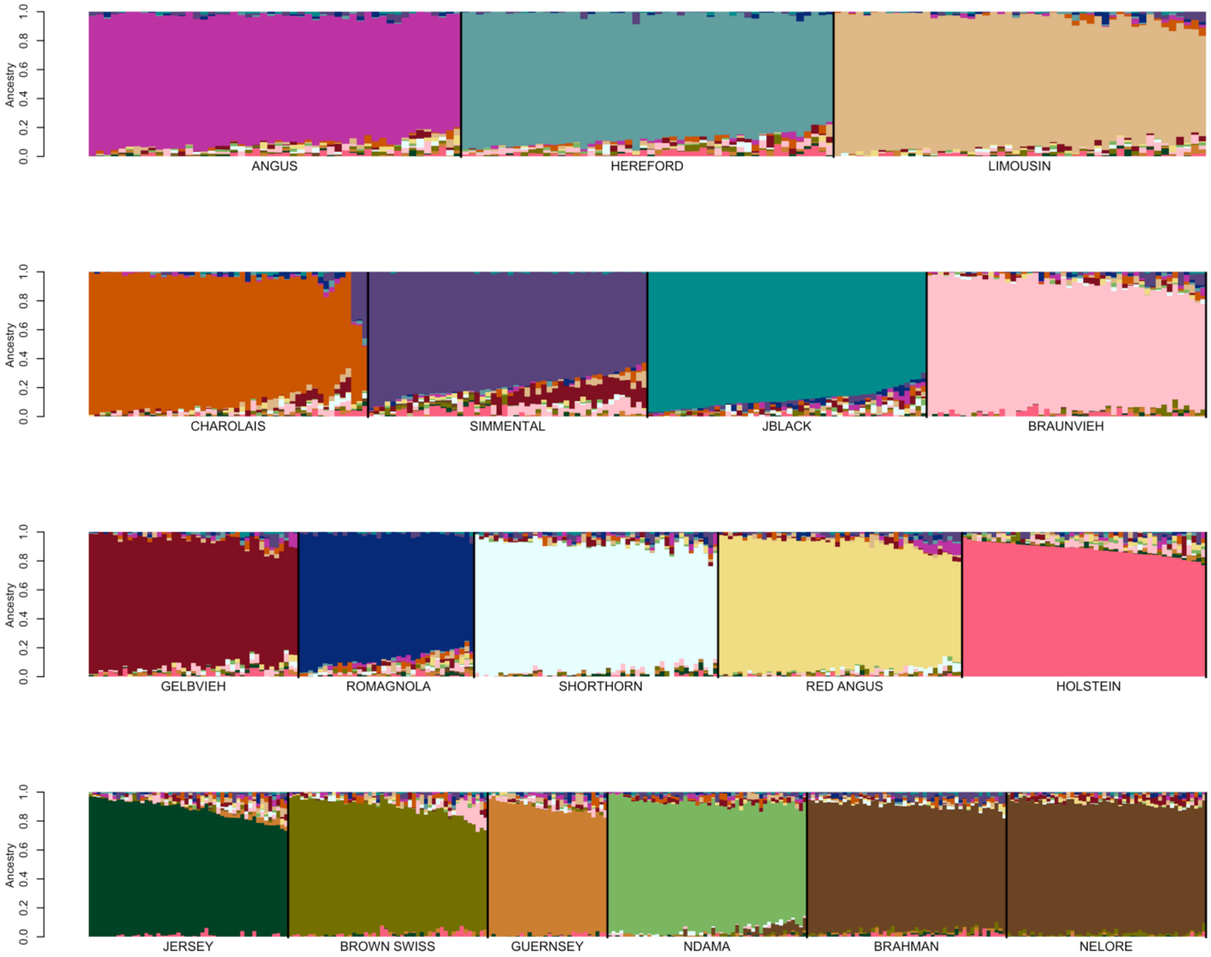


**Fig. S14** SNPweights self-assignment analysis for the reference sample set with ≥75% ancestry to breed of registry and ≤50 individuals per breed using the BC7K marker set..

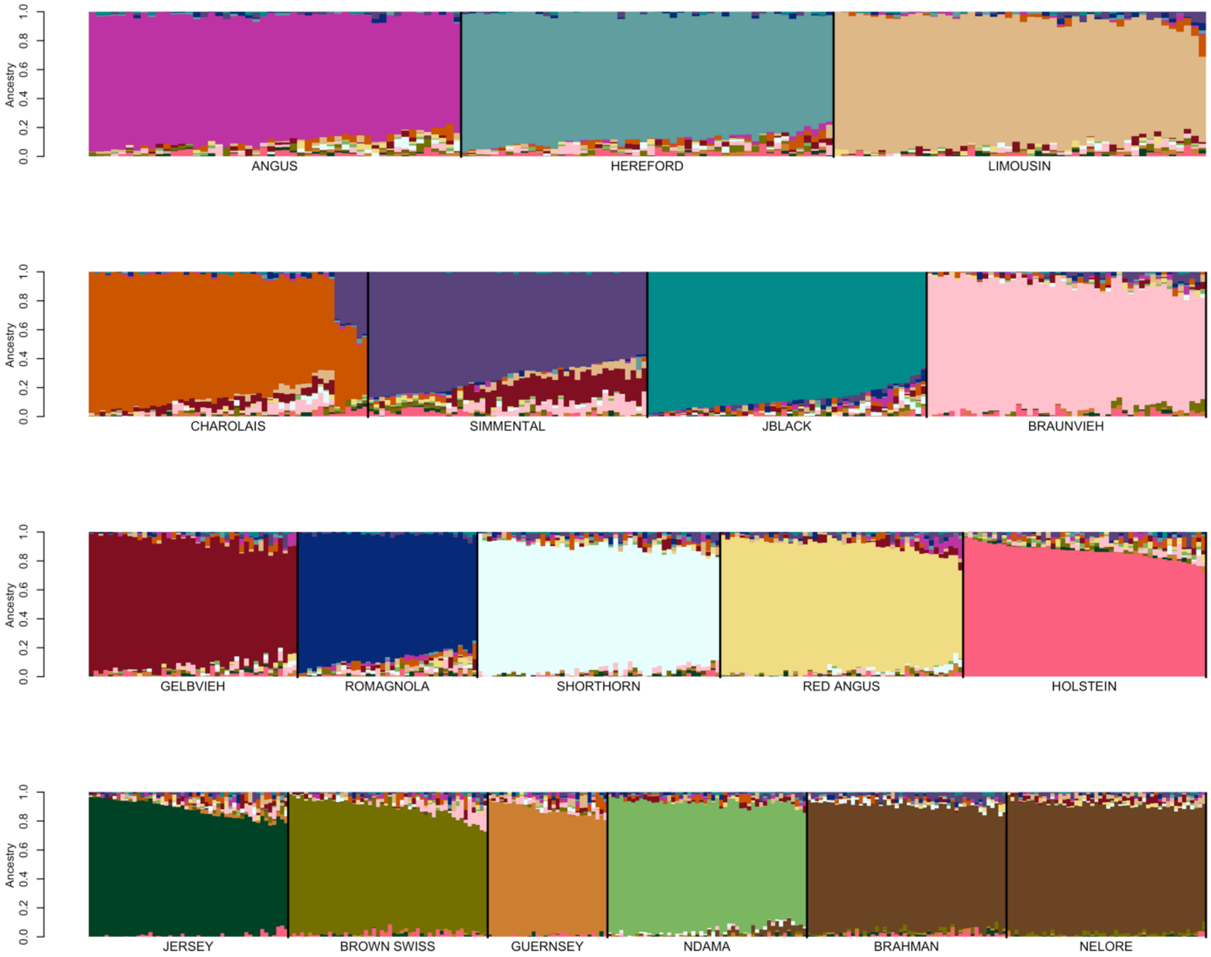


**Fig. S15** SNPweights self-assignment analysis for the reference sample set with ≥70% ancestry to breed of registry and ≤50 individuals per breed using the BC7K marker set.

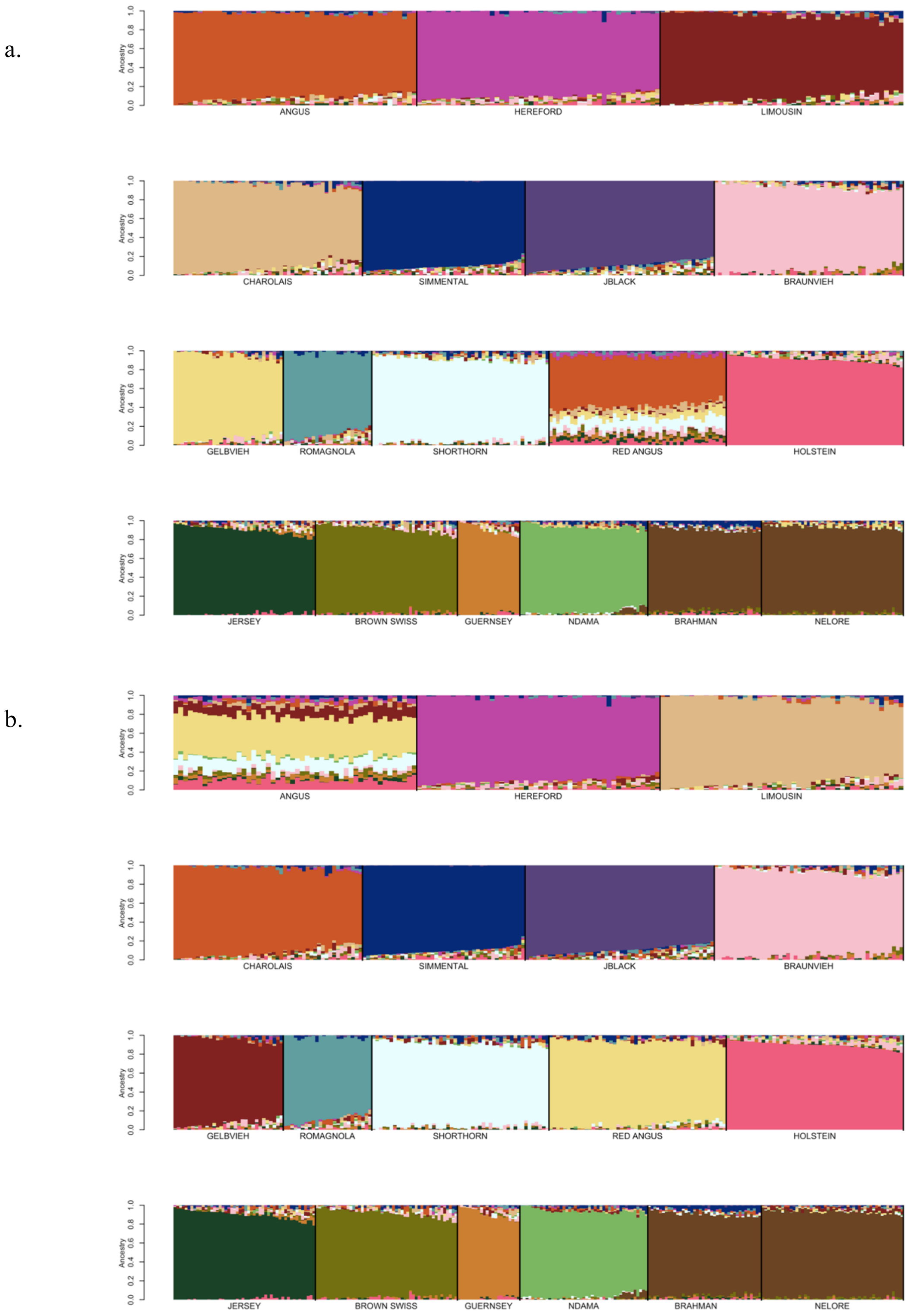


**Fig. S16** SNPweights self-assignment analyses using a reference panel with ≤50 individuals per breed and sampling from the individuals with ≥85% assignment to their breed of registry but with (a) Red Angus or (b) Angus excluded from the reference panel. The Red Angus and Angus individuals in the reference panel were retained for ancestry estimation.
