## Supporting Information for "CRUMBLER: A Tool for the Prediction of Ancestry in Cattle"

**Supplementary Information**

*EIGENSOFT Source Code Edits*

EIGENSOFT packages CONVERTF and SMARTPCA are required for use by SNPweights. However, SMARTPCA within versions of EIGENSOFT beyond 5.0.2 are not compatible with SNPweights. To establish compatibility, the following edits must be made to the SMARTPCA source code:

1. Download EIGENSOFT from https://github.com/DReichLab/EIG/
2. Go to directory /src/eigensrc/
3. Open smartpca.c
4. Find the string: printf("trace: %9.3f\n", y);
5. Remove the comment characters (") to make the code an active line.

In version 7.2.1, the code is located at line #1079. In version 6.1.4, the code is at line #1138. The SMARTPCA program still calculates *trace*, changing the code outputs the value of *trace* for use by SNPweights software.

To recompile the modified code, follow the instructions of the EIGENSOFT authors. This information is found in the README file in the EIGENSOFT download base directory.
