## Supporting Methods for "CRUMBLER: A Tool for the Prediction of Ancestry in Cattle"

**Supplementary Methods.**

**Preliminary fastSTRUCTURE analyses conducted on subsets of breeds.**

*Angus vs Simmental (Fig. S2)*
FastSTRUCTURE analysis was conducted for 5,552 registered Angus and 15,858 registered Simmental. To reduce the potential effects of unequal sample sizes, three analyses were performed. Each analysis included all 5,552 registered Angus and 5,286 of the registered Simmentals. In total, 4,390 Angus were predicted to have at least 97% Angus ancestry in all three analyses and 1,583 Simmental were predicted to have at least 97% Simmental ancestry and were retained for further analysis.

*Angus vs Gelbvieh (Fig. S3)*
FastSTRUCTURE analysis was conducted for 5,552 registered Angus and 12,835 registered Gelbvieh. Three analyses were performed including all 5,552 registered Angus and 4,279, 4,278, and 4,278 registered Gelbvieh, respectively. In total, 4,351 Angus were predicted to have at least 97% Angus ancestry in all three analyses and 6,000 Gelbvieh were predicted to have at least 97% Gelbvieh ancestry and were retained for further analysis.

*Angus vs Limousin (Fig. S4)*
FastSTRUCTURE analysis was conducted for the 4,268 registered Angus identified as having at least 97% Angus ancestry in both the Angus vs Simmental and Angus vs Gelbvieh analyses and 2,734 registered Limousin. In total, 1,470 Angus were predicted to have at least 97% Angus ancestry and 367 Limousin were predicted to have at least 97% Limousin ancestry and were retained for further analysis.

*Angus vs Red Angus (Fig. S5)*
FastSTRUCTURE analysis was conducted for the 4,268 registered Angus identified as having at least 97% Angus ancestry in both the Angus vs Simmental and Angus vs Gelbvieh analyses and 1,377 registered Red Angus. In total, 508 Angus were predicted to have at least 97% Angus ancestry and 124 Red Angus were predicted to have at least 97% Red Angus ancestry and were retained for further analysis.

*Red Angus vs Hereford vs Shorthorn vs Salers (Fig. S6)*
FastSTRUCTURE analysis was conducted for the 1,377 registered Red Angus, 969 registered Hereford, 291 registered Shorthorn, and 68 registered Salers. In total, 700 Red Angus were predicted to have at least 97% Angus ancestry, 335 Hereford were predicted to have at least 97% Hereford ancestry, 212 Shorthorn were predicted to have at least 97% Shorthorn ancestry, and 0 Salers were predicted to have at least 97% Salers ancestry and were retained for further analysis. The Salers was then removed as a reference population.

*Red Angus vs Hereford vs Shorthorn (Fig. S7)*
FastSTRUCTURE analysis was conducted for the 1,377 registered Red Angus, 969 registered Hereford and 291 registered Shorthorn. In total, 518 Red Angus were predicted to have at least 97% Angus ancestry, 348 Hereford were predicted to have at least 97% Hereford ancestry, and

166 Shorthorn were predicted to have at least 97% Shorthorn ancestry, and were retained for further analysis.

*N'Dama vs Nelore vs Brahman (Fig. S8)*
FastSTRUCTURE analysis was conducted for the 98 N'Dama, 941Nelore, and 127 registered Brahman.  In total, 59 N'Dama were predicted to have at least 97% N'Dama ancestry, 86 Brahman were predicted to have at least 97% Brahman ancestry, and 708 Nelore were predicted to have at least 97% Nelore ancestry, and were retained for further analysis.
